# Adapterama I: Universal stubs and primers for 384 unique dual-indexed or 147,456 combinatorially-indexed Illumina libraries (iTru & iNext)

**DOI:** 10.1101/049114

**Authors:** Travis C. Glenn, Roger A. Nilsen, Troy J. Kieran, Jon G. Sanders, Natalia J. Bayona-Vásquez, John W. Finger, Todd W. Pierson, Kerin E. Bentley, Sandra L. Hoffberg, Swarnali Louha, Francisco J. García-De León, Miguel Angel Del Río-Portilla, Kurt D. Reed, Jennifer L. Anderson, Jennifer K. Meece, Samuel E. Aggrey, Romdhane Rekaya, Magdy Alabady, Myriam Bélanger, Kevin Winker, Brant C. Faircloth

## Abstract

Next-generation DNA sequencing (NGS) offers many benefits, but major factors limiting NGS include reducing costs of: 1) start-up (i.e., doing NGS for the first time); 2) buy-in (i.e., getting the smallest possible amount of data from a run); and 3) sample preparation. Reducing sample preparation costs is commonly addressed, but start-up and buy-in costs are rarely addressed. We present dual-indexing systems to address all three of these issues. By breaking the library construction process into universal, re-usable, combinatorial components, we reduce all costs, while increasing the number of samples and the variety of library types that can be combined within runs. We accomplish this by extending the Illumina TruSeq dual-indexing approach to 768 (384 + 384) indexed primers that produce 384 unique dual-indexes or 147,456 (384 × 384) unique combinations. We maintain eight nucleotide indexes, with many that are compatible with Illumina index sequences. We synthesized these indexing primers, purifying them with only standard desalting and placing small aliquots in replicate plates. In qPCR validation tests, 206 of 208 primers tested passed (99% success). We then created hundreds of libraries in various scenarios. Our approach reduces start-up and per-sample costs by requiring only one universal adapter that works with indexed PCR primers to uniquely identify samples. Our approach reduces buy-in costs because: 1) relatively few oligonucleotides are needed to produce a large number of indexed libraries; and 2) the large number of possible primers allows researchers to use unique primer sets for different projects, which facilitates pooling of samples during sequencing. Our libraries make use of standard Illumina sequencing primers and index sequence length and are demultiplexed with standard Illumina software, thereby minimizing customization headaches. In subsequent *Adapterama* papers, we use these same primers with different adapter stubs to construct amplicon and restriction-site associated DNA libraries, but their use can be expanded to any type of library sequenced on Illumina platforms.

## Introduction

Next-generation sequencing (NGS) has transformed the life sciences. The unprecedented amount of sequence data generated by NGS platforms facilitates new approaches, techniques, and discoveries (Ansorge, 2009; Tautz, Ellegren & Weigel, 2010). Reduced costs (Glenn, 2011, 2016) are a major component of NGS success because cost reduction enables many studies that were previously infeasible. Although NGS costs per read have dropped tremendously, the minimum cost to obtain any amount of NGS data (i.e., the minimum buy-in cost) remains high, particularly when researchers want to collect small amounts of DNA sequence data from large numbers of individual samples in a single run. These buy-in costs are largely driven by the money required to purchase adapters containing unique identifying sequences that allow tagging and tracking of samples sequenced in multiplex (Box 1). For example, the purchase price for a subset of 96, single-index, TruSeq-equivalent adapters described in Faircloth & Glenn (2012) would require an initial investment of at least $3,161 (US; $11,321 with TruGrade^®^ purification), and this investment is exclusive of the additional costs to purchase other necessary library preparation reagents and consumables. A second problem for researchers wishing to collect smaller amounts of sequence data from many samples sequenced in multiplex is the relatively limited number of indexed adapters that are available. Although several publications (e.g., Meyer & Kircher, 2010; Faircloth & Glenn, 2012; Rohland & Reich, 2012) and commercial products (e.g., Illumina Nextera, Illumina, San Diego, CA, USA; Bioo Scientific NEXTflex-HT, Bioo Scientific, Austin, TX, USA) provide schemes for indexing hundreds of individuals sequenced in multiplex, most of these approaches do not facilitate individually tagging many thousands of samples at low cost so that samples can be pooled into a single sequencing run. Given the increasing capacity of high-end Illumina instruments (e.g., Illumina NovaSeq), this is a significant and growing issue. A third constraint that has long been known (Kircher, Sawyer & Meyer, 2012) is that Illumina instruments can mismatch the read(s) and index sequence(s) by hopping or swapping indexes (Sinha et al., 2017; Costello et al., 2018), causing sequence misidentification and other problems. Uniquely tagging each index position significantly reduces these problems (Kircher, Sawyer & Meyer, 2012; Illumina, 2017; Costello et al., 2018). As a result, library preparation methods that reduce costs while simultaneously increasing the number of samples that can be tagged and sequenced together would benefit many types of research.

In this first paper of the *Adapterama* series, we present the key components of an integrated system for producing 384 uniquely dual-indexed (or 147,456 combinatorially-indexed) Illumina libraries at low cost (Figs. 1, S1). We build this integrated system on top of previous developments introduced by Illumina (2008) and others (e.g., Meyer & Kircher, 2010; Fisher et al., 2011), and we show that it is possible to significantly reduce library preparation costs by changing from full-length adapters that incorporate tags in the Illumina TruSeq strategy to shorter universal adapter stubs and indexing primers (hereafter referred to as the iTru strategy; which is similar to the original Illumina indexing strategy [Illumina 2008]). Simply moving from a TruSeq indexing strategy to the iTru indexing strategy, while maintaining a single indexing position, can reduce costs by more than 50% (Table 1). When taking advantage of the dual-indexing offered by our iTru strategy, researchers can reduce costs by at least an order of magnitude relative to TruSeq (Table 1). This method is also extensible to the Illumina Nextera adapter sequences (Syed, Grunenwald & Caruccio, 2009; Adey et al., 2010), hereafter referred to as the iNext approach (Figs. S1-S2; File S1). We focus on describing the iTru system because TruSeq is more commonly used than Nextera and to simplify presentation of the system (details of the iNext system are generally given in the supplemental figures and files). In subsequent *Adapterama* manuscripts, we extend the system presented here for a variety of applications (e.g., amplicon sequencing and RADseq), but we use our iTru or iNext indexing primers throughout (Fig. S1).

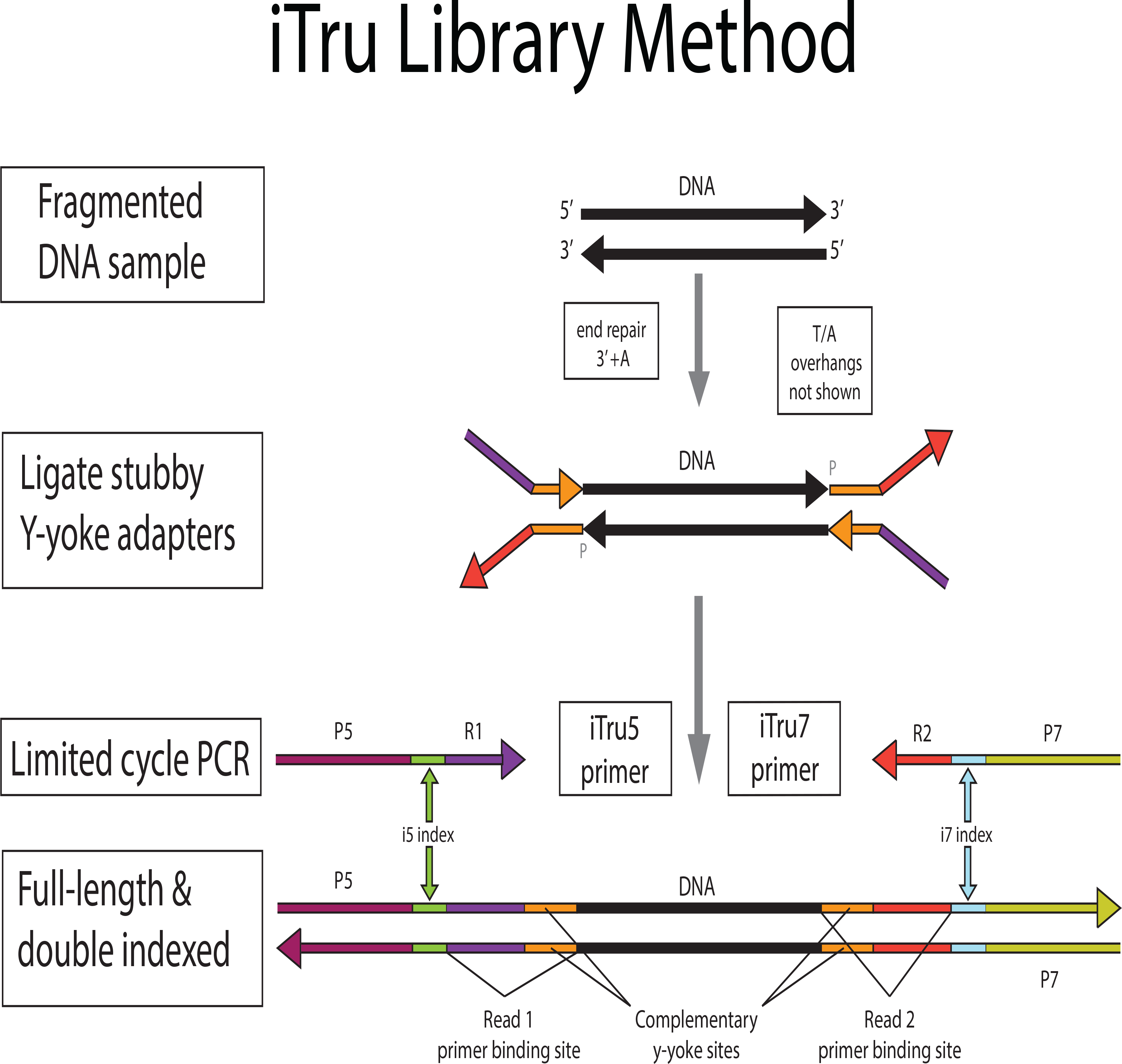

**Table 1:** Comparison of oligonucleotide numbers and costs when using varying numbers of independent tags. Cost estimates assume 2-stage library preparations and list prices from Integrated DNA Technologies, 25 nmole synthesis scale, with oligonucleotides delivered in plates. An index length of 8 nucleotides is used with an edit distance ≥3 for iTru and an edit distance ≥2 for Illumina.

| **Uniquely Indexed Libraries** | **Library Type** | **Index Positions** | **Stub Adapter Oligos** | **Long Adapter Oligos** | **Indexed Primers** | **Adapter Cost + Primer Cost (US \$)** |
| --- | --- | --- | --- | --- | --- | --- |
| 96 | TruSeq* | 1 | 0 | 1+96 | 0 [2<sup>#</sup>] | \$4,019 + \$18 |
| 96 | TruSeq Nano HT | 2 | 0 | 8 + 12 | 0<sup>§</sup> | \$4,560<sup>§</sup> + \$0 |
| 96 | iTru<sup>a</sup> | 1 | 2 | 0 | 1 + 96 | \$45 + \$1,617 |
| 96 | iTru<sup>b</sup> | 2 | 2 | 0 | 8 + 12 | \$45 + \$344 |
| 384 | TruSeq* | 1 | 0 | 1+384 | 0 [2<sup>#</sup>] | \$16,029 + \$18 |
| 384 | iTru<sup>a</sup> | 1 | 2 | 0 | 1 + 384 | \$45 + \$6,416 |
| 384 | iTru<sup>b</sup> | 2 | 2 | 0 | 16+24 | \$45 + \$689 |
| 9216 | TruSeq* | 1 | 0 | 1 + 9216<sup>c</sup> | 0 [2<sup>#</sup>] | \$392,049 + \$18 |
| 9216 | iTru<sup>a</sup> | 1 | 2 | 0 | 1 + 9216<sup>c</sup> | \$45 + \$153,539 |
| 9216 | iTru<sup>b</sup> | 2 | 2 | 0 | 96+96 | \$45 + \$3,333 |
| 74,304 | iTru<sup>b</sup> | 2 | 2 | 0 | 192 + 387 | \$45 + \$9,915 |

\* Original TruSeq approach with custom adapters (cf. Faircloth & Glenn, 2012); kits are no longer available, but the method can be home-brewed (cf. Fisher et al., 2010), or the adapters can be used with reagents from TruSeq Nano kits.
<sup>#</sup> P5 and P7 primers are used.
<sup>§</sup> Price includes all library preparation reagents, not just adapters; P5 and P7 primers are included in kit.
<sup>a</sup> Libraries contain both i5 and i7 tags, but only one iTru5 primer is used for all samples, thus only the i7 tags are informative and are sequenced (cost efficient with old versions of HiSeq $\leq 2500$ kits). This method is no longer recommended, but illustrates cost differences.
<sup>b</sup> Both the i5 and i7 indexes are informative and are sequenced.
<sup>c</sup> Tags of 11 nucleotides are required for 9216 tags of edit distance $\geq 3$ .

Here we outline the ideas underlying genomic library construction for Illumina sequencers, and we provide some historical perspective on Illumina library preparation for researchers new to Illumina sequencing. Following this introduction, we describe our iTru design, which modifies Illumina’s original library construction method and extends the approach to include indexes on both primers (i.e., double-indexing; *c.f.*, Kircher, Sawyer & Meyer, 2012). The iTru method (Figs. 1-3) produces: 1) libraries that are compatible with all Illumina sequencing instruments and reagents; 2) libraries that can be pooled (i.e., multiplexed) with other Illumina libraries; 3) libraries that can be sequenced using standard Illumina sequencing primers and protocols; and 4) data that can be demultiplexed with standard Illumina software packages and pipelines.

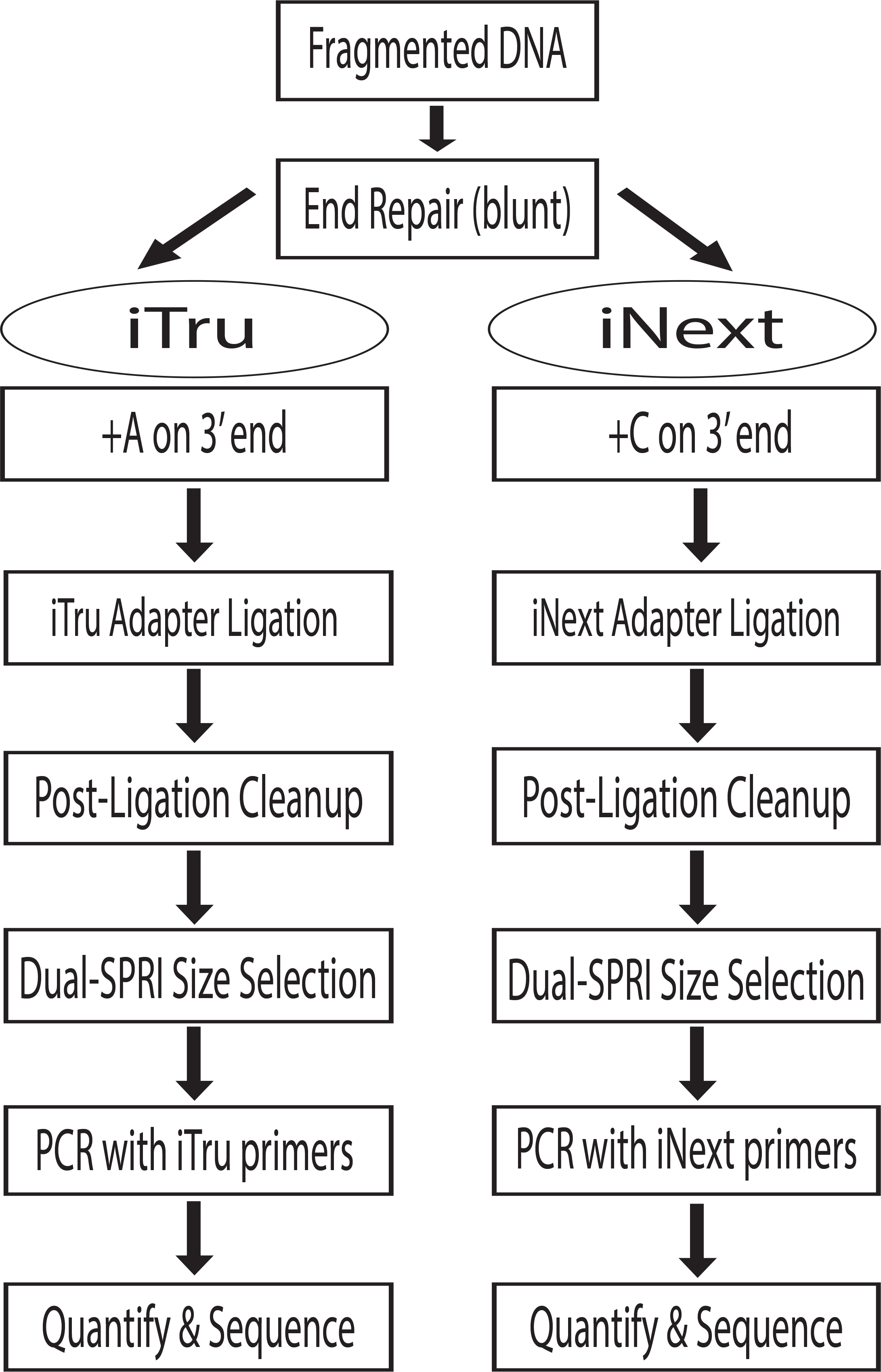

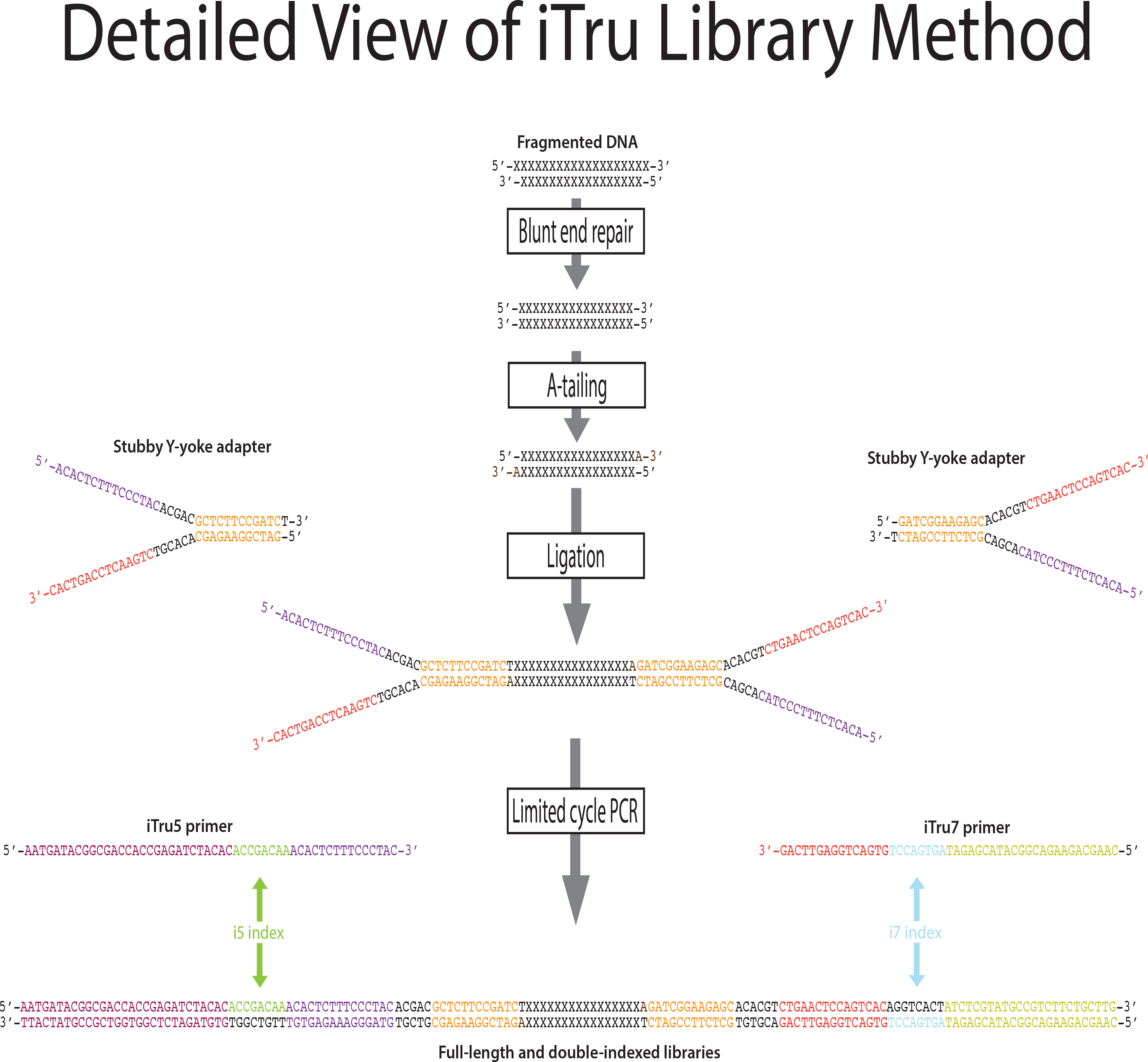

### Illumina libraries

DNA molecules that can be sequenced on Illumina instruments require specific primer-binding sites (i.e., adapters; Box 1) on each end. The procedure to incorporate the adapters to the DNA insert is generally referred to as “library preparation”. Library preparation of genomic DNA, in its most common form, involves randomly shearing DNA to a desired size range (e.g., 200-600 bp); end-repairing and adenylating the sheared DNA; adding synthetic, double-stranded adapters onto each end of the adenylated DNA molecules using T/A ligation; and using limited-cycle PCR amplification to increase the copy number of valid constructs (Figs. 1-3, S3; *c.f.* Fig. S2; Fig. S4).

Illumina library preparations differed from their early competitors (chiefly 454) because their double-stranded adapters used a Y-yoke design to increase library construction efficiency (Bentley et al., 2008; Greigite, 2009). The Y-yoke structure of the adapters allows each starting DNA molecule to serve as two templates, requiring ≥3 cycles of PCR to produce complete double-stranded library molecules (Fig. S3). The DNA molecules resulting from these preparations (Figs. 1-3; Fig. S4) contain: 1) outer primer-binding sites (P5 and P7) used to capture individual DNA molecules on the surface of Illumina flow cells and clonally amplify them; 2) separate primer-binding sites (Read 1 and Read 2), located internal to the P5 and P7 sites, that allow directional sequencing of both DNA strands; and 3) short DNA sequences, known as indexes (Box 1; see below), inserted into the P7 side of the adapter molecule (Illumina, 2008; Fig. 4, i7 index, sequence obtained from Index Read 1; the i5 index was added subsequently, see below).

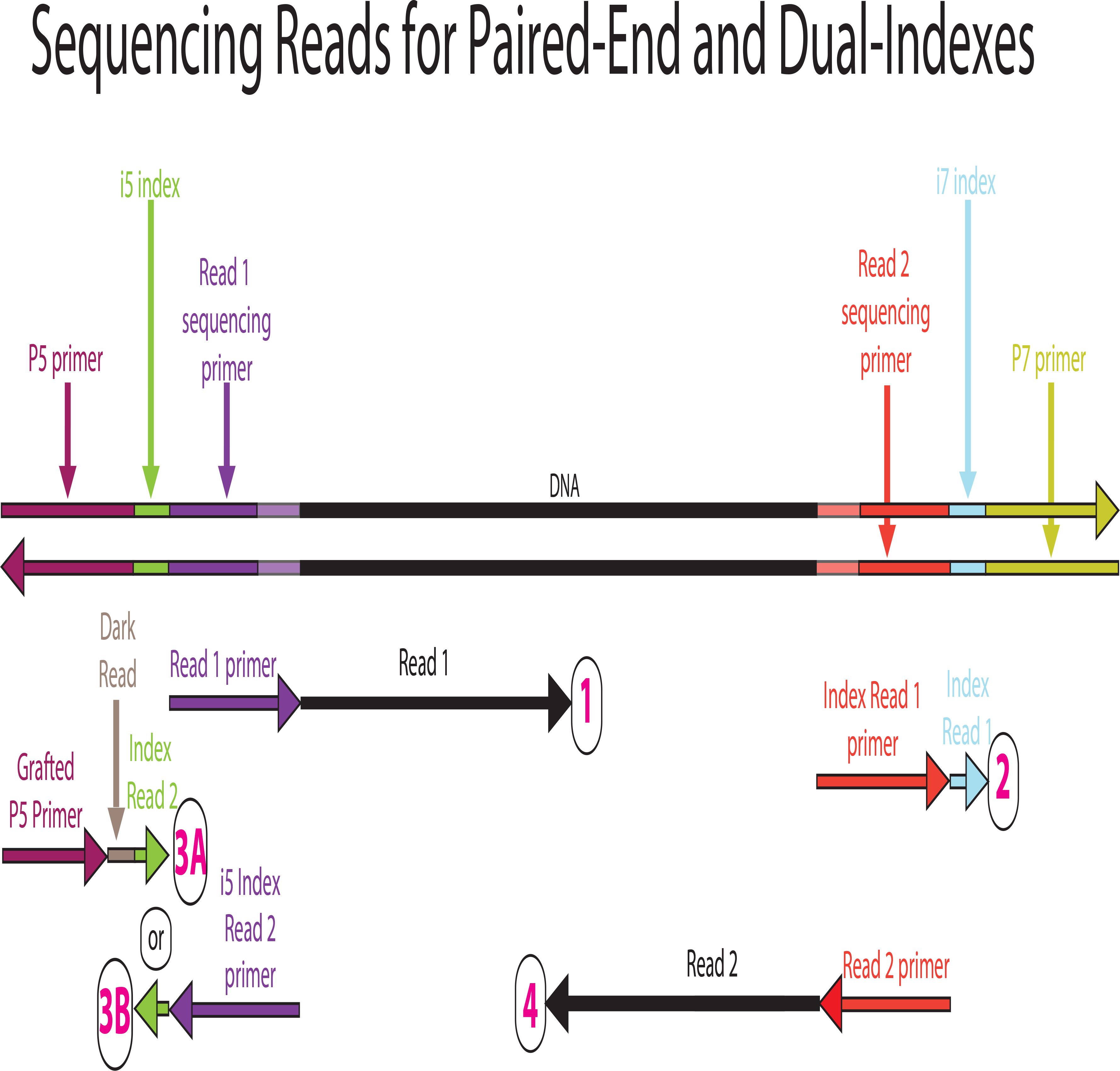

### Indexing

Indexing strategies are generally meant to individually identify different DNA samples by incorporating unique DNA sequences into the library constructs (Shoemaker et al., 1996; Binladen et al., 2007; Glenn et al., 2007; Hoffmann et al., 2007; Meyer et al., 2007; Craig et al., 2008). Indexed libraries can then be pooled together (multiplexed) in a single sequencing lane. During sequencing, individual molecules are captured on the surface of the Illumina flow cells, the individual molecules are clonally amplified, and up to four separate sequencing reactions take place sequentially, each creating a separate sequencing read (Fig. 4). After sequencing, computer software matches the observed index sequence for each molecule to a list of samples with expected indexes (i.e., using a sample sheet; File S2) and parses the bulk data back into its component parts (i.e., demultiplexing, e.g., using bcl2fastq [Illumina, 2017]).

In practice, the history and current status of Illumina indexing strategies is quite complicated (e.g., Illumina, 2018a), with several transitions among different adapter systems that resulted from changing capabilities of sequencing instruments. Illumina originally created 12 different i7 indexes (Fig. 1; Figs. 3-4) to allow pooling of up to 12 samples, and the company later increased the number of i7 indexes for certain applications to 48. The original Illumina i7 indexes had a length of six nucleotides (nt) and were constructed such that ≥2 substitution errors were needed to turn one index into another—an effort to minimize sample confusion as a result of sequencing error. Sequencing errors on Illumina instruments are primarily substitutions; thus, Illumina’s initial indexes were designed to be robust to substitution sequencing errors. Deletions, however, are the primary errors of oligonucleotide synthesis (i.e., synthesis of the adapters and/or primers used to make the indexed libraries). It is, therefore, desirable to have indexes that are robust to insertions and deletions (indels) as well as substitutions, thus conforming to an edit-distance metric and limiting the assignment of sequences to the wrong sample (Faircloth & Glenn, 2012). When index sets have edit-distances ≥3, then error correction can be employed, but this distance criterion is frequently violated (Faircloth & Glenn, 2012).

Building upon earlier in-house and external efforts, Illumina introduced a product (Nextera kits) that used an i5 index and an i7 index (i.e., dual-indexing; see Box 1, Fig. 1, and below) each of which were longer (8 nt) and, at that time, conformed to the edit-distance metric. Nextera adapters use the same sequences for interaction with the flow-cell (i.e., P5 and P7; Fig. 1), but have unique Read 1 and Read 2 sequences relative to TruSeq (Fig. S2; Fig. S4). Thus, Illumina does not recommend combining Nextera and TruSeq libraries within a single sequencing lane (Illumina, 2012; but see below). Illumina subsequently incorporated 8 nt, dual indexes into the TruSeq system with their release of TruSeqHT. Although the Illumina TruSeqHT indexes are robust to insertion, deletion, and substitution errors, the updated TruSeqHT i7 indexes do not maintain an edit-distance ≥3, when compared to other TruSeq HT i7 indexes in the same set or when combined with all previous Illumina i7 indexes, and so do not allow proper error correction (Fig. S5; File S3). Regardless, the TruSeqHT indexing system is more robust, accurate, and flexible than previous approaches, and researchers can index template DNA molecules using the i7 indexes alone (single-indexing) or in combination with i5 indexes (dual-indexing).

Dual indexing on the Illumina platform means that indexes can be used combinatorially (Kircher, Sawyer & Meyer, 2012; Faircloth & Glenn, 2012). Major advantages of the dual-indexing strategy include: 1) the need for fewer oligonucleotides to index the same number of samples in multiplex (e.g., 8 + 12 = 20 primers produce 8 × 12 = 96 unique tag combinations); 2) concomitantly reducing the cost of production, inventory, and quality control (QC) (i.e., it is less expensive to produce, maintain stocks of, and do QC on 20 primers than 96); and 3) the universality of the approach—dual-indexing is compatible with both full-length adapters (e.g., TruSeqHT libraries) or universal adapter stubs and primers (e.g., Nextera, iNext, or iTru). As noted above, combinatorial indexing is susceptible to index hopping which results in sequences being assigned to the incorrect samples, whereas using unique sequences at multiple index positions (e.g., unique dual-indexes) significantly reduces these problems (Kircher, Sawyer, & Meyer, 2012; Illumina, 2017; Costello et al., 2018).

### Illumina-compatible libraries

Illumina’s libraries have been the industry’s gold standard for sequence quality on Illumina platforms, but their library preparation kits are among the most expensive available. The number of indexes offered by Illumina was limited to ≤48 and the number of dual-index combinations ≤96, until subsequent releases of additional indexes for the Nextera system, which can dual-index up to 384 samples (Illumina, 2018b). Most recently, Illumina has partnered with Integrated DNA Technologies, Inc. (IDT, Coralville, IA, USA) to develop a set of 192 (96 + 96) indexed adapters that also contain unique molecular identifiers (https://support.illumina.com/downloads/idt-illumina-truseq-ud-indexes-sample-sheet-templates.html; UMIs, Box1) to improve multiplexing, mitigate sample misassignment due to index hopping, and detect PCR duplicates (IDT, 2018; MacConaill et al., 2018). Alternative commercial kits have been produced to increase efficiency, reduce GC bias (Aird et al., 2011; Kozerewa et al., 2009), and/or increase the number of indexes, but costs remain high and the total number of commercially available indexes still generally remains ≤384.

A variety of library preparation methods have also been described by research groups that reduce per-sample costs relative to most commercial kits (e.g., Meyer & Kircher, 2010 [MK-2010]; Fisher et al., 2011 [F-2011]; see Head et al., 2014 for others). The MK-2010 and F-2011 methods are in widespread use, but they do have some shortcomings. For example, the MK-2010 method: 1) specifies HPLC purification of adapter oligonucleotides, which increases start-up costs dramatically and can lead to contamination from previous oligonucleotides that were purified on the same HPLC columns; 2) relies on hairpin suppression of molecules with identical adapter ends (instead of using a Y-yoke adapter) which is efficient with smaller inserts (e.g., <200 bp) but loses efficiency with increasing insert length; and 3) relies on blunt-ended ligation, which allows the formation of chimeric inserts. The F-2011 method introduced the idea of “on-bead” library preparation, which increases efficiency and reduces costs; thus, many commercial kits have subsequently incorporated similar on-bead library preparation approaches. Limitations of the F-2011 method include use of: 1) custom NEB reagents, not in the standard catalog or available in small quantities; 2) large volumes of enzymes; and 3) Illumina adapters and primers, which increase costs and limit the number of samples that can be pooled.

Our approach builds upon many of the previous approaches introduced by Illumina, MK-2010, F-2011, Rohland & Reich (2012), and others to develop library preparation methods for genomic DNA that overcome many of these limitations. We describe adapters, primers, and library construction methods that produce DNA molecules equivalent to and compatible with Illumina’s TruSeqHT libraries (and, separately, Nextera libraries, see File S1; Table 2). Our method extends the number of available index combinations from 96 × 96 to 384 × 384, while maintaining a minimum edit-distance of ≥3 between all indexes. We demonstrate the effectiveness of our combinatorial indexing primers by controlled quantitative PCR experiments, and we demonstrate the utility of our system by preparing and sequencing iTru libraries from organisms with varying genome size and DNA quality.

**Table 2:** Comparison of Nextera, iNext, iTru, and TruSeq Nano HT library preparation methods.

| **Library Type** | **Nextera** | **iNext** | **iTru** | **TruSeq Nano HT** |
| --- | --- | --- | --- | --- |
| **Input DNA (ng)** | Intact ($\geq 50$) | Sheared ($\geq 100^{\#}$) | Sheared ($\geq 100^{\#}$) | Sheared ($\geq 100$) |
| **Repair ends** | N/A | Yes | Yes | Yes |
| **Add DNA overhang** | N/A | C | A | A |
| **Ligate adapter** | Tagmentation | iNext stub | iTru stub | TruSeq |
| **Limited cycle PCR primers** | Nextera or iNext* | Nextera or iNext | iTru | P5 and P7 |
| **Advantages** | Least time | Lower cost, high diversity | Lower cost, high diversity | Industry standard |
| **Disadvantages** | Higher cost, lower diversity, less randomness<sup>§</sup> | More prep. time than Nextera | More prep. time than Nextera | Higher cost, more input DNA, more prep. time; not for sequence capture |

\* Note, iNext primers are not specified as biotinylated, and thus will not work interchangeably with Nextera libraries that use streptavidin beads to capture/normalize/purify libraries unless biotins are added. Using unmodified iNext primers requires other purification and normalization procedures.
<sup>§</sup> Tagmentation does not insert adapters into the genome as randomly as shearing the DNA.
<sup>#</sup> Hyper Prep Plus Kits (KapaBioSciences) allow input as low as 1 ng of intact DNA.

## Materials & Methods

### Adapter and primer design

We modified the Illumina TruSeq system by dividing the adapter components into two parts: 1) a universal Y-yoke adapter “stub” that comprises parts of the Read 1 and Read 2 primer binding sites plus the Y-yoke; and 2) a set of amplification primers (iTru5, iTru7), parts of which are complementary to the Y-yoke stub and which also contain custom sequence tag(s) for sample indexing (Fig. 1; Fig. 3; Table 3; File S4) as well as the sequences (P5, P7) necessary for clonal amplification on Illumina flow cells. The iTru Y-yoke adapter has a single 5’ thymidine (T) overhang and can be used in standard library preparations that produce insert DNA with single 3’ adenosine (A) overhangs. We designed a large set of indexed amplification primers (iTru5, iTru7; File S4) that contain a subset of our custom 8 nt sequence tags (from Faircloth & Glenn, 2012), as well as an initial set that incorporated the TruSeq HT indexes (i.e., D5xx for iTru5 and D7xx for iTru7) which could serve as controls. All iTru5 indexes are compatible with Illumina indexes. Some of the iTru7 indexes are not compatible with Illumina indexes (i.e., edit-distance is ≤2). We grouped the iTru primers with our sequence tags into clearly identifiable, numbered sets (100 and 300 series) that are compatible with 8 nt indexes in the standard Illumina TruSeqHT primers, as well as Illumina v2 8 nt indexes (including the 6 nt indexes converted to 8 nt via addition of invariant bases from the adapter). We also created several additional numbered sets (200 and 400 series) of iTru primers that are compatible with all other primers and sequence tags in our iTru system, but which are not compatible with all Illumina indexes. We then balanced the base composition of all iTru primers in all numbered sets in groups of eight for iTru5 and groups of 12 for iTru7, because balanced base composition is critical for successful index sequencing (Illumina, 2016; see Discussion for additional information on combining small numbers of libraries).

**Table 3:**
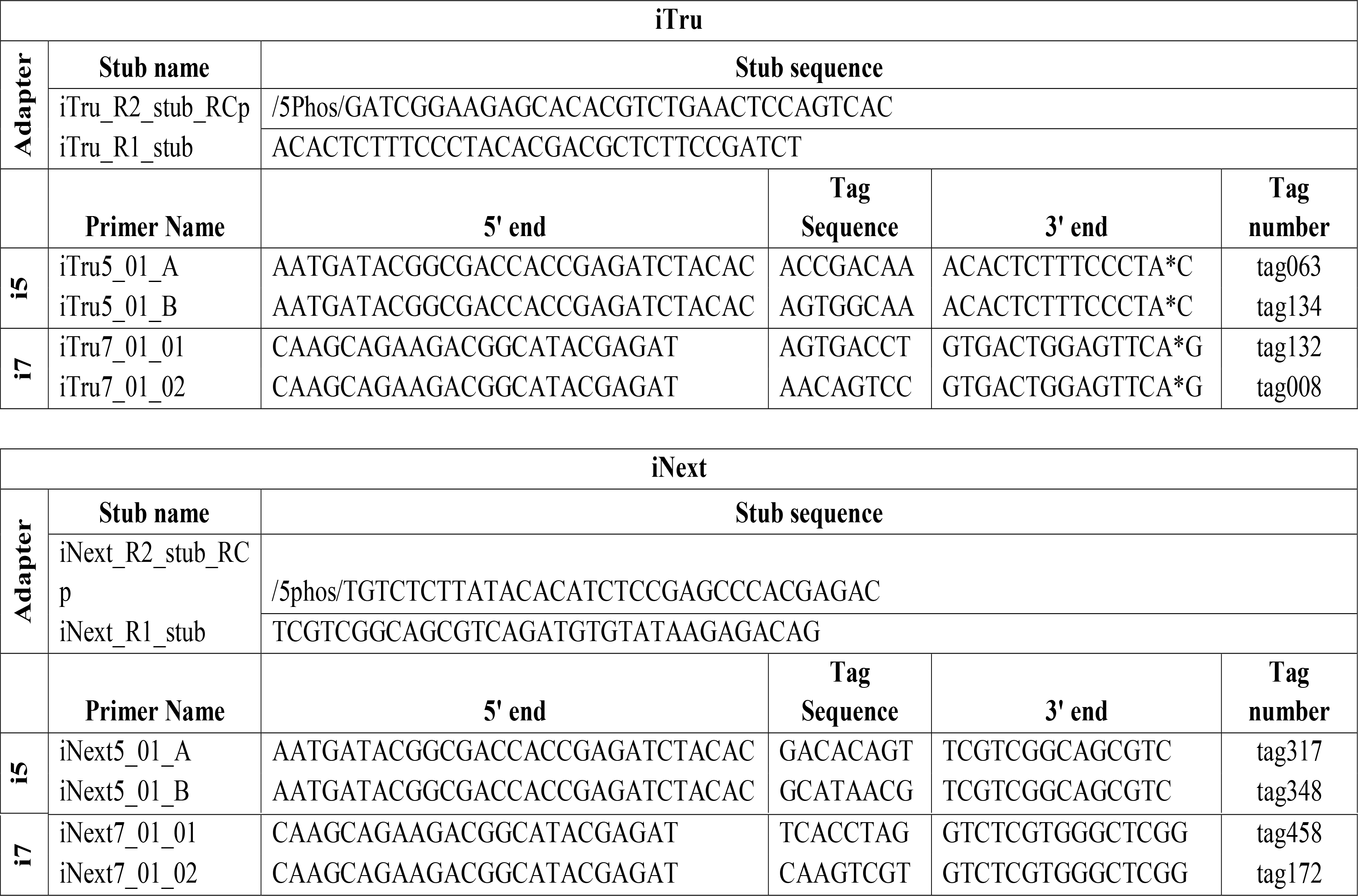
iTru and iNext adapter stub oligonucleotides and tagged primer sequences. All sequences are given in 5’ to 3’ orientation. To make it clear which portions are constant among all tagged primers, as well as to identify function, the tagged primers are given in three pieces (the invariant 5’ end, the tag sequence which varies among primers, and the invariant 3’end), but the primers are obtained as a single contiguous fusion of these three pieces. Complete balanced sets of primers are available as Supplemental Files (4, 15). Adapter stub oligonucleotides must be hydrated and annealed prior to use (Supplemental File 7).

We ordered the components of our Y-yoke adapter stubs and iTru primers from IDT, with standard desalting purification only. We modified the adapter stub sequence by phosphorylating the 5’ end of iTru_R2_stub_RCp oligonucleotide (Fig. 1; Table 3), and we modified each of the iTru primer sequences by adding a phosphorothioate bond (Eckstein, 1985) before the 3’ nucleotide of each sequence to inhibit degradation due to the exonuclease activity of proof-reading polymerases (Skerra, 1992), which are commonly used in library preparation. Following initial small-scale orders, we ordered sets of iTru primers, placing the iTru5 and iTru7 primers into every other column (iTru5) or row (iTru7) of 96-well plates, with 0.625 or 1.25 nmol aliquots in replicate plates (Files S4-S5). We hydrated newly synthesized primers to 10 µM in the plate and 5 µM prior to use (File S6). Subsequently, we ordered the complete set of 384 iTru5 and 384 iTru7 primers in 96-well plates with 1.25 nmol aliquots (Files S4-S5).

### Validation of iTru primers by quantitative PCR (qPCR)

To determine whether our indexed iTru5 and iTru7 primers were biasing amplification, we selected a subset of iTru7 (n=160) and iTru5 (n=48) primers for qPCR validation. To validate the iTru primers, we prepared a pool of adapter-ligated chicken DNA using an inexpensive, double-digest RAD approach (3RAD; Graham et al., 2015, Bayona-Vásquez et al., 2019) that produces a DNA construct having 5’ and 3’ ends identical to our Y-yoke adapter. We then set up quantitative PCR reactions with 5 µL GoTaq qPCR Master Mix (Promega, Madison, WI, USA), 1 µL each forward and reverse primer at 5 µM, 2 µL adapter-ligated DNA at 0.12 ng/µL, and 1 µL H_2_O. Working under the assumption that Illumina primers have been validated as unbiased by Illumina, we tested all forward (iTru5) primers with Illumina D701 as the reverse primer, and we tested all reverse (iTru7) primers with Illumina D501 as the forward primer. We ran all primer tests in duplicate on an Applied Biosystems StepOnePlus (Thermo Fisher Scientific, Waltham, MA, USA) using the following conditions: 95°C for 2 min, then 40 cycles of 95°C for 15 s, and 60°C for 1 min. Because we needed to run multiple plates of qPCR to test all of the primers, we included the iTru5 set 2 primer A (iTru5_02_A) and the iTru7 set 2 primer 1 (iTru7_02_01) on all plates to provide a baseline reference for iTru5 or iTru7 primer performance. We determined the threshold cycle (C_T_) using the default settings of the StepOnePlus, we averaged C_T_ values from replicate runs, and we calculated Delta C_T_ for each iTru primer using two approaches. First, we evaluated the relative performance of all iTru5 and iTru7 primers by subtracting the C_T_ of the iTru5 or iTru7 primer being tested from the average C_T_ of all iTru5 or iTru7 primers. Second, we evaluated the performance of all iTru5 and iTru7 primers by subtracting the baseline reference C_T_ of iTru5_02_A from the C_T_ of the iTru5 primer being tested and by subtracting the baseline reference C_T_ of iTru7_02_01 from the C_T_ of the iTru7 primer being tested. We expected that unbiased primers would not deviate from the average and/or baseline performance by more than 1.5 PCR cycles (>1.5 C_T_), a value that should encompass the stochasticity seen between independent PCR reactions as a result of small, unavoidable primer concentration and other amplification performance differences.

### Implementation in *E. coli* and eukaryote libraries: DNA source

To test the performance of both our Y-yoke adapters and the iTru system in a variety of library preparation scenarios, we prepared genomic libraries from DNA of various types and quality. As a simple, known source of control DNA, we used *Escherichia coli* k-12 strain MG1655 (hereafter *E. coli*; Roche, Basel, Switzerland), which has a high-quality genome sequence available (GenBank accession NC_000913; 4.6 Mb) and is commonly used for quality control of sequencing libraries. To examine how our iTru system performed with DNA of varying quality and complexity, we also prepared iTru libraries from DNA that we isolated from six samples from a diverse array of species (two sharks, one tarantula, one jellyfish, and a coral). We isolated each of these DNA sources using a variety of techniques commonly used in many labs, including commercial kits, salting out, or CTAB Phenol-Chloroform extraction (Table 4; also see File S1 for additional details about testing iNext). These samples represent the range of species, sampling conditions, and DNA isolation techniques that are commonly encountered in model and non-model organism studies, and the taxa we sampled included particularly challenging specimens (i.e., tarantula, coral and jellyfish) that have previously performed poorly with commercial library preparation kits. Before library preparation, we fragmented *E. coli* genomic DNA to 400-600 bp using a Covaris S2 (Covaris, Woburn, MA, USA), and we fragmented genomic DNA (normalized to 23 ng/µL) to 400-600 using the Bioruptor UCD-300 sonication device (Diagenode, Denville, NJ, USA).

**Table 4:** Results from initial iTru library preparation and sequencing tests of DNA from sharks and challenging non-model organisms. The Illumina i7 index sequences where used in these tests. Protocol 1: EZNA Tissue DNA KIT (Omega Bio-Tek, USA); Protocol 2: Aljanabi & Martínez (1997); Protocol 3: CTAB-Phenol.

| Sample ID | Common Name | Species | DNA Extraction Method | i7 Index ID | Raw Index Count | Number of Read Pairs | Primary Objective | Usable Reads | putative mtDNA contig size in bp (mean coverage) | Microsats Identified<sup>2</sup> |
| --- | --- | --- | --- | --- | --- | --- | --- | --- | --- | --- |
| MaF 5 | white shark | *Carcharodon carcharias* | Protocol 1 | 705 | 1,930,539 | 1,805,638 | mtDNA | 1,722,562 | 17,103 (46x)<sup>3</sup> | - |
| MaF 19 | white shark | *Carcharodon carcharias* | Protocol 2 | 707 | 2,075,236 | 1,927,792 | mtDNA | 2,003,858 | 17,138 (31x)<sup>3</sup> | - |
| MaF 10 | silky shark | *Carcharhinus falciformis* | Protocol 1 | 706 | 1,438,468 | 1,358,550 | mtDNA | 1,800,534 | 17,285 (22x)<sup>4</sup> | - |
| MaF 1 | Tarantula | *Brachypelma vagans* | Protocol 1 | 701 | 985,171 | 934,406 | msats | 80,790 | - | 563 |
| MaF 16 | cannonball jellyfish | *Stomolophus spp.* | Protocol 3 | 703 | 959,516 | 909,401 | msats | 591,608 | - | 92,668 |
| MaF 9 | Coral | *Poritespanamensis* | Protocol 1 | 702 | 3,449,711 | 3,298,155 | msats | 1,549,718 | 18,628 (50x)<sup>5</sup> | 7.322 |
| Total |  |  |  |  | 10,838,641 | 10,233,942 |  |  |  |  |

<sup>1</sup> Only includes high quality reads with inserts of 250 bases; excluded reads generally due to short insert length due to degraded input DNA.
<sup>2</sup> Identified using default parameters in PAL-finder (Castoe et al., 2012).
<sup>3</sup> Díaz-Jaimes et al. (2016)
<sup>4</sup> Galván-Tirado et al. (2016)
<sup>5</sup> Del Rio-Portilla et al. (2016)

### Implementation in *E. coli* and eukaryote libraries: library construction

Prior to library preparation, we annealed the iTru adapter sequences to form double-stranded, Y-yoke adapters by mixing equal volumes of the iTru_R1_stub and iTru_R2_stub_RCp oligos at 100 µM, supplementing the mixture with 100 mM NaCl, heating the solution to 98°C for 2 min in a thermal cycler, and allowing the thermal cycler to slowly cool the mixture to room temperature (File S7).

We prepared genomic iTru libraries from *E. coli* using kits, reagents, and protocols from Kapa Biosystems (Roche, Basel, Switzerland), with minor modifications to the manufacturer’s instructions. The major change we made was to ligate the universal iTru adapter stubs (Table 3; File S4) to the 3’-adenylated (i.e., +A) DNA fragments, and then use the iTru5 and iTru7 primers with TruSeqHT indexes for limited-cycle amplification (Figs. 1-3). For the eukaryotic libraries, we further modified the manufacturer’s instructions by using half-volume reaction sizes with the following two changes. We used an inexpensive alternative to commercial SPRI reagents (Sera-Mag SpeedBeads, Thermo-Scientific, Waltham, MA, USA; see File S8) in all cleanup steps. After adapter ligation, we performed a post-ligation cleanup followed by SPRI dual-size selection using first 0.55x PEG/NaCl and then an additional 0.16x SpeedBeads which also contains PEG/NaCl. We outline step-by-step methods for this approach in File S9.

### Sequencing

We quantified libraries using a Qubit 2.0 Fluorometer (Thermo Fisher Scientific, Waltham, MA, USA) and KAPA qPCR, checked for index diversity (File S10), and then normalized and pooled all libraries at 10 nM (File S11). We also ensured the quality of library pools by running 1 µL on a Bioanalyzer High Sensitivity chip (Agilent Technologies, Santa Clara, CA, USA). We combined the iTru and iNext *E. coli* library pools (File S1) with samples from other experiments, and we sequenced the combined pools using a single run in Illumina MiSeq v2 500 cycle kit (PE250). We combined the eukaryotic libraries with additional TruSeq libraries from other experiments and sequenced these on a separate run of Illumina MiSeq v2 500 cycle kit to produce PE250 reads.

### Sequence analysis

After sequencing, we demultiplexed reads using Illumina software (bcl2fastq v 1.8 – 2.17; Illumina 2013). We then imported reads to Geneious 6.1.7 – R9.0.4 and trimmed adapters and low-quality bases (<Q20). We removed reads with inserts of <125 bases prior to all downstream analyses. We mapped *E. coli* reads back to NC_000913 using the Geneious mapper (fastest setting, single iteration). We assembled reads from the eukaryotic libraries using the Geneious assembler (fastest setting), and we extracted contigs of 250 to 450 bp from eukaryotic libraries of tarantula, jellyfish, and coral for downstream microsatellite searches using msatCommander 1.0.8 (Faircloth, 2008). We also used PAL_FINDER v0.02.03 (Castoe et al., 2012) to enumerate microsatellites within read-pairs that had inserts ≥250 bases. Finally, we extracted contigs of approximately 17 kb from the shark libraries, and we used MEGA-BLAST searches to determine which of these contigs represented shark mtDNA genomes (Díaz-Jaimes et al., 2016). We did the same with approximately 18 kb fragments from the coral (Del Rio-Portilla et al., 2016).

### Larger-scale tests

Following initial validation of the iTru primers and the utility of the iTru library preparation approach, we placed the iTru system into an extensive test phase in which we routinely used this approach for library construction within our own labs while we also made all components of the iTru system available to dozens of other labs. To demonstrate the utility of our approach across a variety of projects, we analyzed read count data from four of these studies (n=576 libraries) that used the iTru system as part of a workflow for target enrichment of ultraconserved elements (UCEs; Faircloth et al., 2012). These included 90 iTru libraries prepared by our group from cichlid fishes (McGee et al., 2016), 183 iTru libraries prepared by a second group from carangimorph fishes (Harrington et al., 2016), 100 iTru libraries prepared by a third group from ants (Faircloth et al. 2015; Blaimer et al., 2016), and 203 iTru libraries prepared by our group from birds. For the bird libraries, we prepared one batch of standard Illumina libraries (n=10) and 2 batches of iTru libraries (n=203), which allowed us to look at sample-to-sample differences in read counts returned from standard Illumina libraries relative to our iTru libraries. One of the two batches of iTru libraries (n=92) combined standard Illumina primers (D5xx; which we used on *E. coli*) on the P5 side with iTru7 primers on the P7 side. The second batch (n=111) combined iTru5 primers on the P5 side with iTru7 primers on the P7 side. The first batch allowed us to assess iTru7 performance separate from that of iTru5, while the iTru5+iTru7 libraries allowed us to assess performance of the full iTru system relative to all other combinations. For all remaining libraries within the other projects, each group followed the protocols for iTru library preparation described above using combinations of only iTru5 and iTru7 primers.

Following library preparation and PCR amplification, each laboratory combined all libraries into equimolar pools containing 8-12 libraries and followed a standardized protocol for target enrichment of UCE loci (http://ultraconserved.org; Faircloth et al. 2012). After enrichment, each group used a Bioanalyzer to determine the insert size of enriched libraries and, to reduce the variance in number of reads sequenced from each pool, quantified pools using a commercially available KAPA qPCR kit. Prior to sequencing, all research groups used the average fragment size distribution and qPCR concentration of each pool to produce an equimolar, project-specific pool-of-pooled-libraries for sequencing with a final concentration of 10 nM. We sequenced the enriched cichlid and carangimorph libraries using different, partial runs of PE150 sequencing on an Illumina NextSeq, the ant libraries using one lane of PE125 sequencing on an Illumina HiSeq 2500, and the bird libraries using two lanes of PE150 sequencing on an Illumina HiSeq 1500 (Rapid Run Mode). For the carangimorph fish libraries, we wanted each sample to receive 0.5% of the total number of reads in the NextSeq run. For all other libraries, we wanted each library to receive 1% of the total number of reads. After sequencing, we computed the average number of raw reads returned per sample, the 95% confidence interval (95 CI) of reads returned per sample, and the percentage of reads returned per sample.

## Results

### Validation of iTru primers by quantitative PCR (qPCR)

Almost all iTru primers (158/160 iTru7 and 48/48 iTru5) had average C_T_ values within 1.5 cycles of both the average Δ C_T_ and the baseline Δ C_T_ (Fig. S8; File S12), suggesting that our iTru indexed amplification primers amplify successfully (98.7% success for iTru7; 100% success for iTru5) and perform similarly to one another. There were two iTru7 primers that failed to amplify during their initial tests, iTru7_401_07 and iTru7_209_04. We rehydrated a new plate of primers and retested iTru7_401_07, which amplified normally (C_T_ = 19.4, Δ C_T_ (average) = −0.7; Δ C_T_ (baseline) = 1.1) during the retest.

### *E. coli* iTru libraries

The iTru libraries we prepared from *E. coli* returned similar numbers of reads from each iTru library, averaging 973,008 reads per sample (95 CI: 161,044; Fig S9; File S13). Each library contained >400,000 high quality reads that covered >99.99% of the known *E. coli* genome sequence. These results suggest that our genomic iTru library preparation process produces valid constructs for Illumina sequencing, and that iTru dual-indexed libraries pooled at equimolar ratios return roughly similar amounts of sequence data (Fig. S9), although we combined libraries at equimolar ratios prior to sequencing using fluorometry which can result in some variation around the targeted read number for each library.

### Eukaryote iTru libraries

We successfully sequenced all eukaryotic genomic libraries prepared using the iTru system and the libraries returned an average of 1,806,440 reads per sample (95 CI: 743,337; Table 4). Using a genome skimming approach, we sequenced the mitogenomes of the shark and coral samples to an average coverage of 33x and 50x, respectively. We used the contig assemblies from our tarantula, jellyfish, and coral samples to design primers pairs targeting >100 microsatellite loci in each taxon. Although the variance in the number of sequencing reads returned per library was higher among these samples than the *E. coli* libraries, these results demonstrate that the iTru system can be used to prepare libraries from DNA of different organisms extracted using different purification approaches, including DNA that produced very poor results with commercial kits (data not shown).

### Larger-scale tests

Our beta test allowed us to collect sequence data from many different iTru5 and iTru7 primers used to index a variety of iTru libraries from fishes, ants, and birds. Few of the libraries that we or others prepared using the iTru system showed large differences in the desired number of reads sequenced when compared to libraries having Illumina-only adapters/index sequences when viewed in aggregate (Fig. S10) or on an index-by-index basis across projects (Figs. S11-S14; File S14). The iTru primer combinations that sometimes returned a lower number of reads for a particular library in a particular project did not show this behavior in other studies (e.g., compare iTru7_402_07 in Fig. S13 versus Fig. S14), suggesting that the reduction in read numbers results from particular library preparation, pooling, enrichment, and quantification practices for specific samples (i.e., specific experimental errors, library preparation methods, or sample-index interactions) rather than inherently bad iTru indexes/primers.

## Discussion

Our results show that the iTru universal adapter stubs and iTru primers can be used to produce genomic libraries for a variety of purposes. The low variance in C_T_ values among iTru5 and iTru7 primers demonstrates that the different index sequences have minimal effect on the libraries, and our results from real-world tests demonstrate that the iTru system works well with DNA from different extraction methods and of differing quality, quantity, and copy number. The results we present from DNA libraries prepared using the iTru system in our and others’ laboratories show that the approach easily scales to hundreds of libraries prepared, pooled, and sequenced in a single lane, ultimately producing information consistent with the variety of Illumina library techniques we have employed to obtain similar data (e.g., Crawford et al., 2012; McCormack et al., 2013; Smith et al., 2014).

After testing the iTru system in several labs, we made several changes in our approach. The most significant of these were: 1) use a naming scheme that allows researchers to easily identify sets of iTru7 primers that are compatible or incompatible with TruSeq indexes; and 2) to increase the amount of iTru5 and iTru7 aliquoted into plates after oligo synthesis (from 0.625 nmol to 1.25 nmol), which reduced library amplification failures that resulted from improper hydration of low-quantity primers in specific wells of plates. The naming scheme and concentrations used in all supplemental files and the naming scheme we used in the Methods section reflect these changes to minimize confusion. After making these changes, we and others have successfully produced libraries and sequencing reads from all iTru5 and iTru7 primers, libraries for many of the primers are detailed in the supplemental files, and we have no evidence suggesting that any of the primer sequences will not work correctly. The original sets of iTru7 primers (sets 00 – 13) synthesized for beta testing have mixed compatibility with Illumina indexes, thus we encourage beta users to exhaust old stocks and adopt the new sets.

It is important to note that the iTru5 and iTru7 primers are grouped into “balanced” sets of 8 or 12 to minimize problems of index base diversity during sequencing. Index balance problems arise because of the way Illumina platforms detect bases during the sequencing run (Illumina, 2016), and the main issues associated with unbalanced base composition are experienced when relatively few samples are sequenced or when a small number of libraries with unbalanced sequence tags take up a large fraction of the sequencing run. We modeled the original four color-scheme used in HiSeq and MiSeq instruments. Using an entire group of eight iTru5 and 12 iTru7 indexed primers within a sequencing pool where each library is present in equal proportion ensures balanced base representation during the index sequence read(s). We also empirically validated this in the two-channel system used in NextSeq, MiniSeq and NovaSeq platforms. Generally, when researchers multiplex more than one group of eight iTru5 or 12 iTru7 indexed primers, base diversity is even more balanced, although it is always a good idea to check the balance of sequencing tags in all sequencing runs (i.e., use File S10). When less than a whole set of primers (i.e., <8 iTru5 primers or <12 iTru7 primers) are used, or if very few libraries will dominate the percentage of reads within a run, it becomes critical to ensure the tags are sufficiently diverse (i.e., use File S10, which includes separate calculations of base diversity for both color schemes). It is also possible to use the stub ligation products from one sample for multiple PCR reactions with different iTru5, iTru7 primers, or even to pool iTru5 and iTru7 primers, thus creating increased numbers of indexes in a pool from a limited number of samples.

All of the iTru oligonucleotides make use of a single phosphorothioate bond between the penultimate and 3’ base. Phosphorothioate linkages protect the 3’ end of oligonucleotides from some forms of nuclease activity (Ekstein, 1985; Skerra, 1992) such as those introduced by some DNA ligases and polymerases (exonuclease activity is a common contaminant of ligases and an intrinsic activity of proofreading polymerases), but phosphorothioate linkages add a modest cost to each primer (~$3 USD per phosphorothioate linkage). Phosphorothioate linkages are also chiral, so only 50% of synthetic molecules receive protection per linkage, while the other 50% remain susceptible to nuclease activity (Eckstein, 1985). Adding a second phosphorothioate bond can reduce the proportion of unprotected molecules by 50% (thus 75% would be protected and 25% would remain susceptible). Illumina and other vendors often include three or more phosphorothioate linkages at the 3’ end of their oligonucleotides to ensure that a large fraction of the molecules are protected from nuclease activity. We include only a single phosphorothioate linkage in our iTru oligo designs because if we lose the 3’ base, we would rather lose the rest of the molecule instead of rescuing the remaining part of it, which may not function appropriately. This strategy also reduces costs associated with synthesizing the oligonucleotides, although others may prefer to incorporate additional phosphorothioate linkages (e.g., two phosphorotioate linkages would lead to 50% fully protected oligonucleotides and 25% that only lose a single 3’ base).

### Who should adopt this method?

Today, there is great need to efficiently minimize cost per sample by scaling and increasing multiplexing flexibility, especially with the advent of platforms like the NovaSeq 6000 that can yield up to 3000 Gb in a single run. Researchers who need higher capacity to multiplex their Illumina library preparations or who have not yet invested heavily in any other method will likely find our approach attractive. It has a low cost of entry and significant flexibility (see below). The more types of libraries, projects, and samples researchers use, the quicker they will recoup the cost of switching and see savings. Additionally, researchers using MK-2010 to construct libraries with inserts >200 bp, particularly those inserts ≥500 bp, are likely to benefit from using a Y-yoke adapter. Our dual-indexed iTru/iNext libraries also reduce concerns over misassignment because, although index-switching occurs with low probability at both ends of sequences in a library, it rarely affects both ends of the same fragment (Larsson et al., 2018).

Researchers already invested in and using other methods with good success, such as the MK-2010 or F-2011 approaches, may wonder if it is worthwhile to switch. We suggest that it would be reasonable to continue using the MK-2010 and/or F-2011 methods if these are already being used successfully; for these labs, we simply provide some alternative adapters and primers that could be used once existing stocks of MK-2010 and/or F-2011 adapters and primers are exhausted or when new projects requiring unique or larger numbers of uniquely tagged samples are encountered.

### iNext

In addition to the iTru adapters and primers we designed and tested, we have developed a universal adapter stub and sets of primers (iNext; Supplementary File 1) that are compatible with the Illumina Nextera system and the original 8 × 12 Nextera indexes, though they are not compatible with all of the subsequent Nextera indexes. As noted in the methods, both iNext and iTru make use of slightly different subsets of the tags identified by Faircloth & Glenn (2012), and the indexed primer sets and numbering approaches are independent between iNext and iTru (e.g., iNext5_01_A does not have the same sequence tag as iTru5_01_A). Thus, researchers should use the tag sequence or tag number from Faircloth & Glenn (2012) or the tag sequences themselves to determine which indexes are equivalent (e.g., iNext7_07_06 uses tag 113 [AGCTAAGC] as does iTru7_203_10; these should not be combined into a single sequencing pool). Although we demonstrate it is possible to combine iNext and iTru libraries within the same MiSeq run (File S13; the iNext and iTru *E. coli* data come from a single MiSeq run) and have subsequently added iNext or Nextera libraries in limited quantities to several of our iTru library pools run on the MiSeq, we are skeptical that other researchers should or will do this routinely. If researchers want to combine iNext and iTru libraries on a regular basis, it would be worthwhile to run additional experiments and to screen and sort the tags to compile sets with numbering that is consistent, thus facilitating pooling between the two systems.

### Troubleshooting

Although all researchers endeavor to conduct mistake-free experiments, foul-ups are certain to occur. In addition to simple record-keeping errors, a very common mistake is flipping the orientation of one of the strip tubes containing iTru primer aliquots. Thus, it is critical to have the capacity to quickly and easily determine what index sequences and combinations are present within a sequencing run. We have developed a small and fast python program (Supplementary File S15) that can count the indexes within a file of reads that were not assigned to specific samples during demultiplexing (i.e., the undetermined reads from bcl2fastq).

### Other applications and future modifications

It is possible to use the iTru system for a variety purposes beyond what we describe here. For example, we have used the iTru system for making RNAseq libraries using KAPA library kits, as well as NEB Ultra II and Ultra II FS (New England Biolabs, Ipswich, MA, USA). Nearly any approach that yields double-stranded template molecules with a single adenosine can be used with no significant modifications to what we have described. One of the attractive features of our system is that it separates the primers and stubs into more manageable units. In other *Adapterama* papers, we use these same iTru primers with different adapter stubs to construct double-to quadruple-indexed amplicon libraries (Glenn et al., 2019), double-digest restriction-site associated DNA (3RAD; Bayona-Vásquez et al., 2019), and RADcap (Hoffberg et al., 2016) libraries. All of these extensions facilitate library preparation, sequencing, and bioinformatic processing of these types of data while also significantly reducing costs.

Having separate primers and adapter stubs simplifies and reduces costs associated with modification or swapping out of the universal Y-yoke adapters (Table 3; Files S4; S16), creating opportunities for further research and protocol development. For example, if researchers wanted to optimize library preparation for low levels of input DNA, then implementing an adapter stub in a stem-loop configuration [*e.g.,* NEB Next Ultra; (New England Biolabs, Ipswich, MA, USA)] would be worth investigating. Similarly, adapters containing uracils that are broken at the uracil sites by USER (NEB M5505) or uracil-DNA-glycosylase (UDG; e.g., NEB M0280) plus APE 1 (e.g., NEB M0282) facilitate a variety of designs with potentially beneficial characteristics worth exploring, especially for mate-pair libraries. However, given recent advances in commercial kits that reduce buffer exchanges and increase efficiency (e.g., KAPA Hyper and HyperPlus and NEB Ultra II and UltraII FS, which require as little as 1 ng of input DNA), it is likely that the use of such high efficiency approaches combined with the iTru adapters and primers will be sufficient for the vast majority of applications where samples derive from ≥1000 eukaryotic cells.

## Conclusions

We describe an approach that uses a single universal adapter stub and relatively few PCR primers to produce many Illumina libraries. The approach allows multiple researchers to have unique primer sets so that libraries from individual researchers can be pooled without worrying about tag overlap. These primers can also be used with a variety of other application-specific adapters described in subsequent *Adapterama* papers for amplicon and RADseq libraries (Bayona-Vásquez et al., 2019; Glenn et al., 2019; Hoffberg et al., 2016). By modularizing library construction, researchers are free to focus on the development of new application-specific tags. Taking advantage of the many available tags also creates opportunities for low-cost experimental optimization attempts. Although the adapters and primers we describe are specific to Illumina, many of the ideas can easily be extended to Ion Torrent, Pacific Biosystems, Oxford Nanopore, and other sequencing platforms (Glenn et al., 2007).

## Acknowledgements

Oligonucleotide sequences © 2007-2019 Illumina, Inc. All rights reserved. Derivative works created by Illumina customers are authorized for use with Illumina instruments and products only. All other uses are strictly prohibited. We thank Rigoberto Delgado Vega, Yann Henaut, Salima Machkour M’Rabet, Fausto Valenzuela, Eduardo Balart, David Paz, Carolina Galvan, Liza Gomez Daglio, and Pindaro Díaz Jaimes for providing samples, Erin Lipp for generously sharing laboratory space and equipment, Rahat Desai, Megan Beaudry, Julia Frederick and Will Thompson for helpful edits, and our colleagues at the Georgia Genomics and Bioinformatics Core and the Georgia Advanced Computing Resource Center. We thank Richard Harrington, Matt Friedman, and Thomas Near (carangimorph fishes), and Michael Branstetter, John Longino, and Phil Ward (ants) for allowing us to use read-count information from their respective studies. Finally, we acknowledge and thank Lisa Ortuno (deceased) for her enthusiastic support of this work; our world was enriched while she shared it with us.

