## Supplementary material for "Adapterama I: Universal stubs and primers for 384 unique dual-indexed or 147,456 combinatorially-indexed Illumina libraries (iTru & iNext)": Fig. S1

### Double Index 2-Step Library Method

Input DNA sample

1) Attach application-specific stubs

2) Use limited cycle PCR to add indexes and extend adapters

Full-length & double indexed

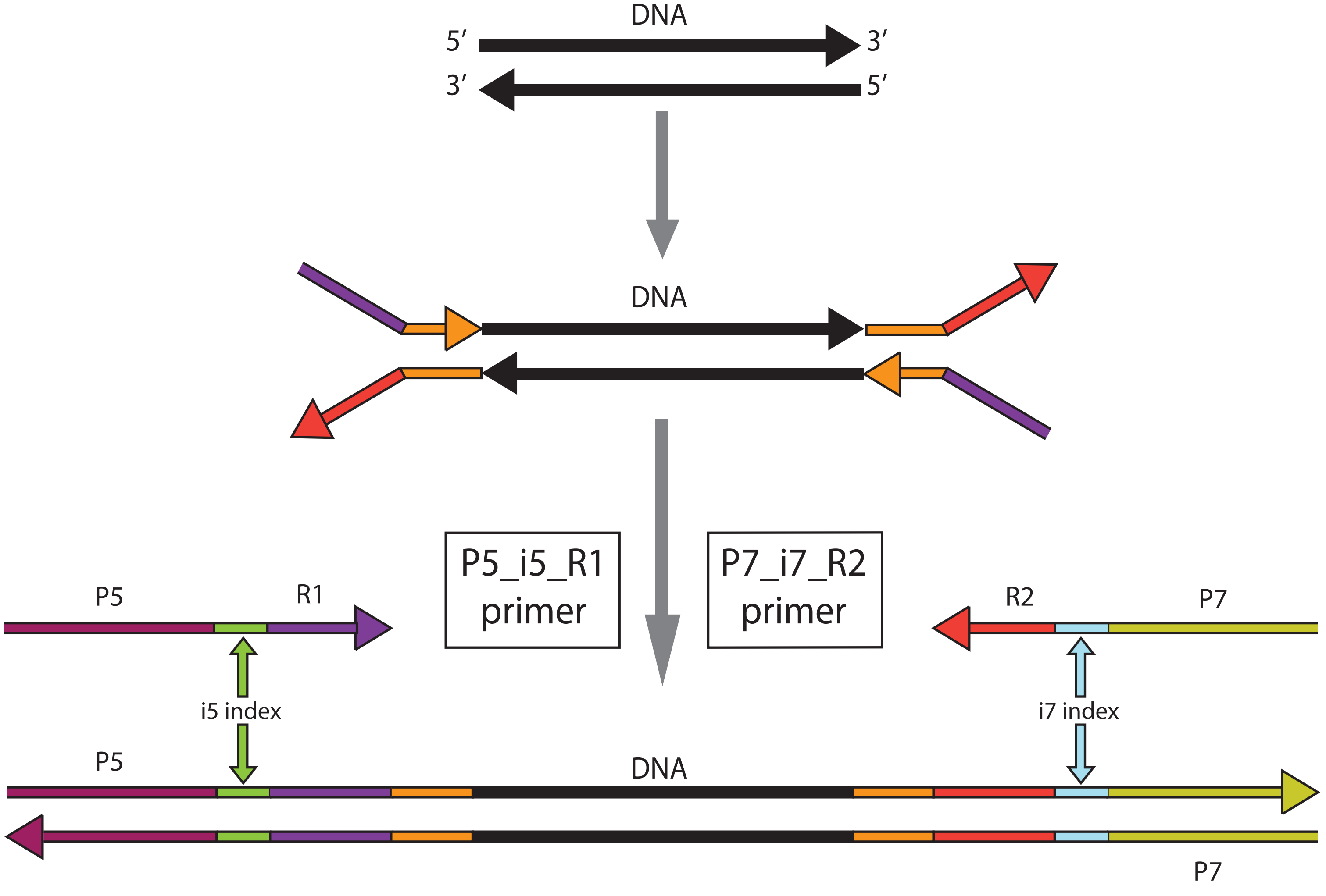
