## Supplementary figures and images for "Adapterama I: Universal stubs and primers for 384 unique dual-indexed or 147,456 combinatorially-indexed Illumina libraries (iTru & iNext)"

### Fig. S2

#### iNext Library Method

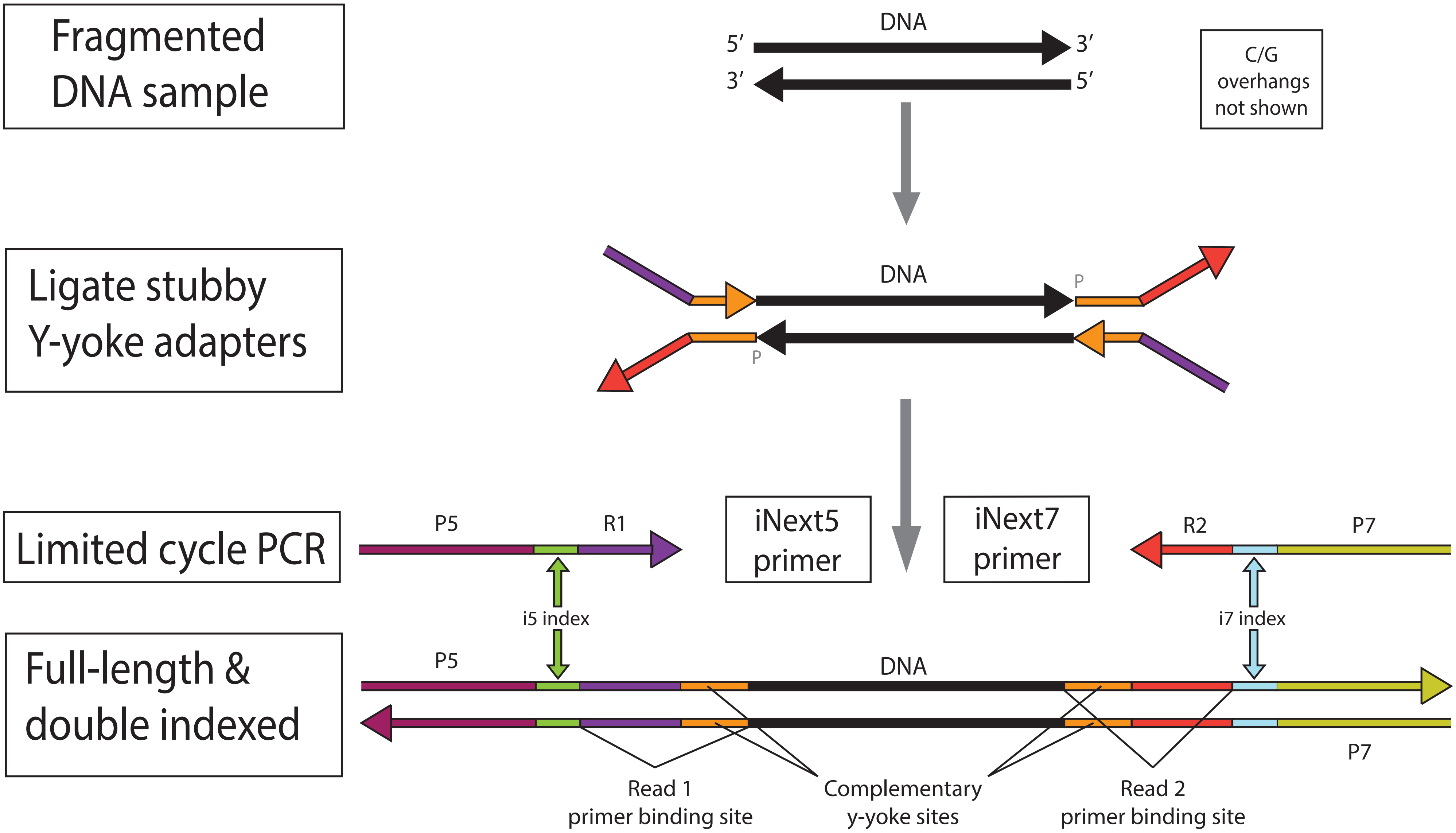

### Fig. S3

#### Addition of Double-Index with PCR

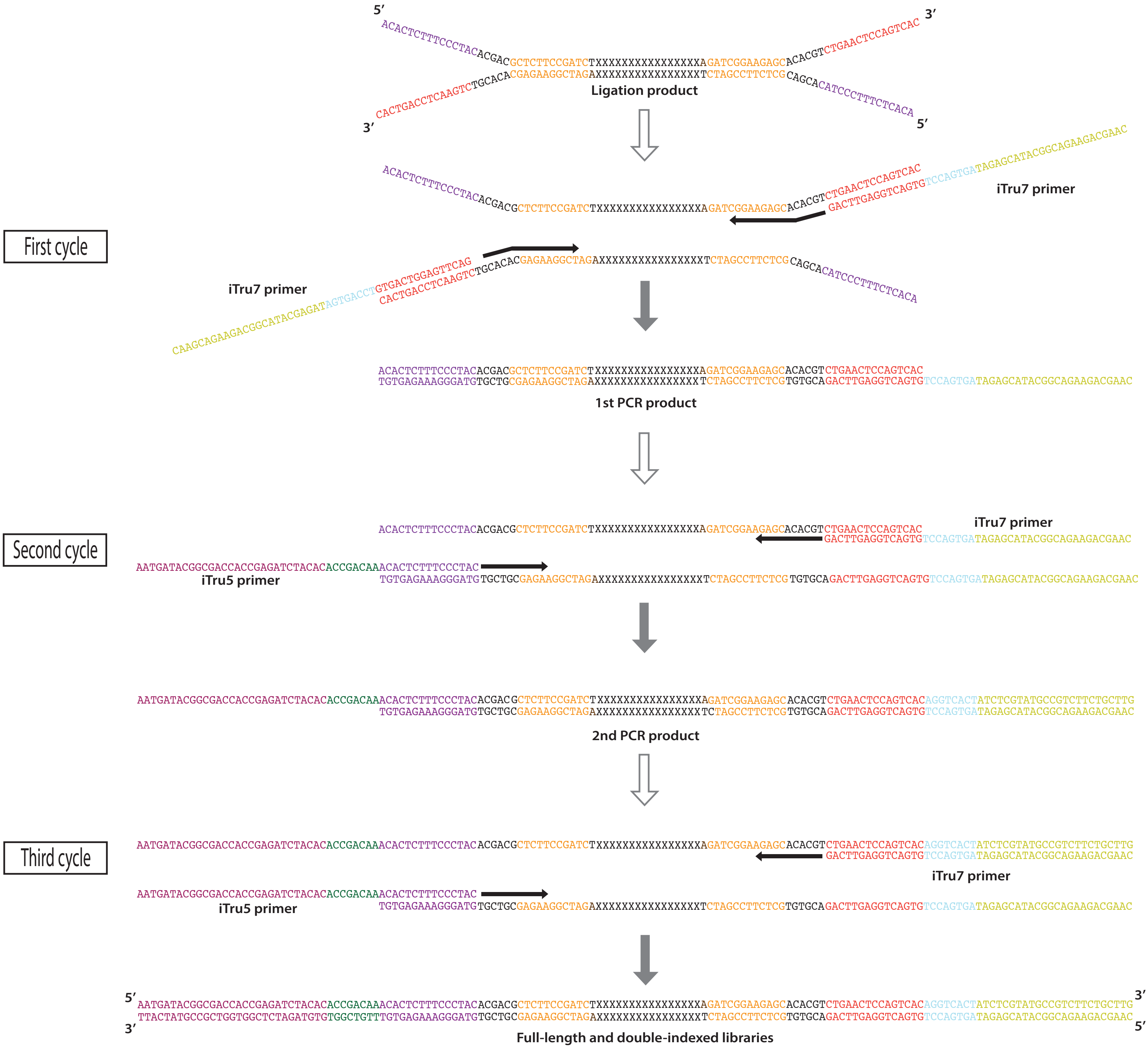

### Fig. S4

#### Complete Library Molecule TruSeq (iTru) vs. Nextera (iNext)

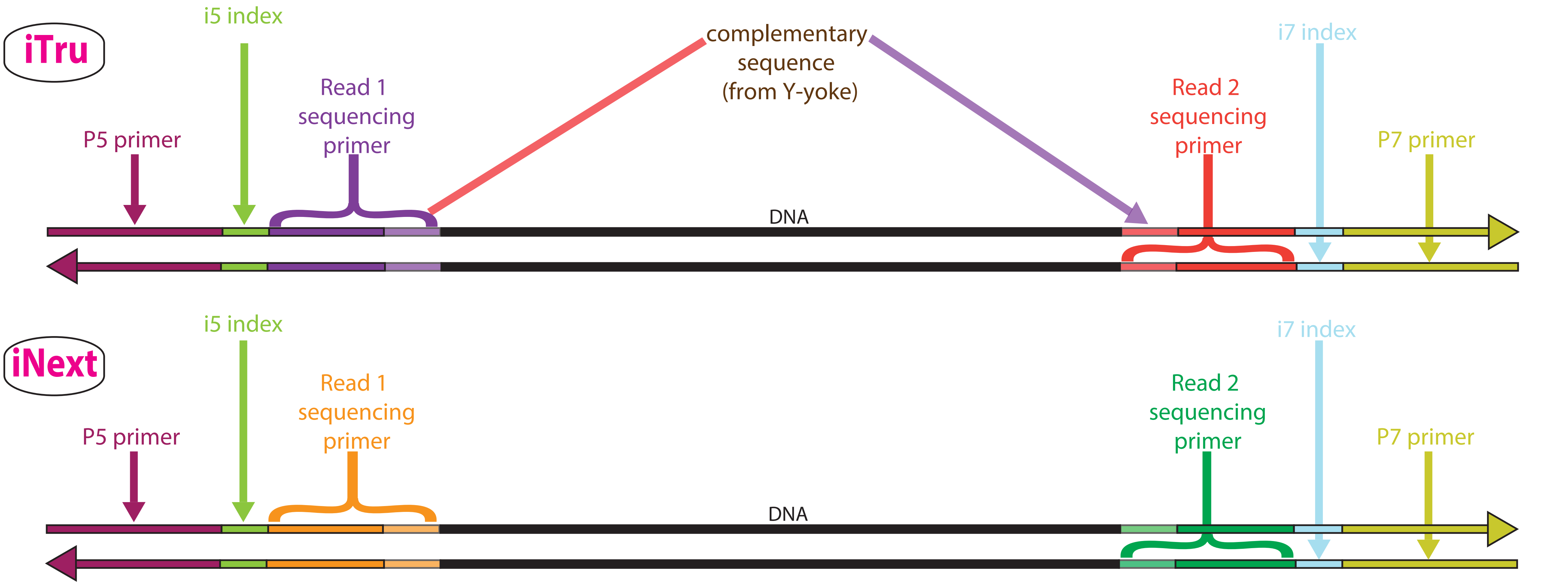

### Fig. S6

#### Complete Library Molecule and Sequencing Reads

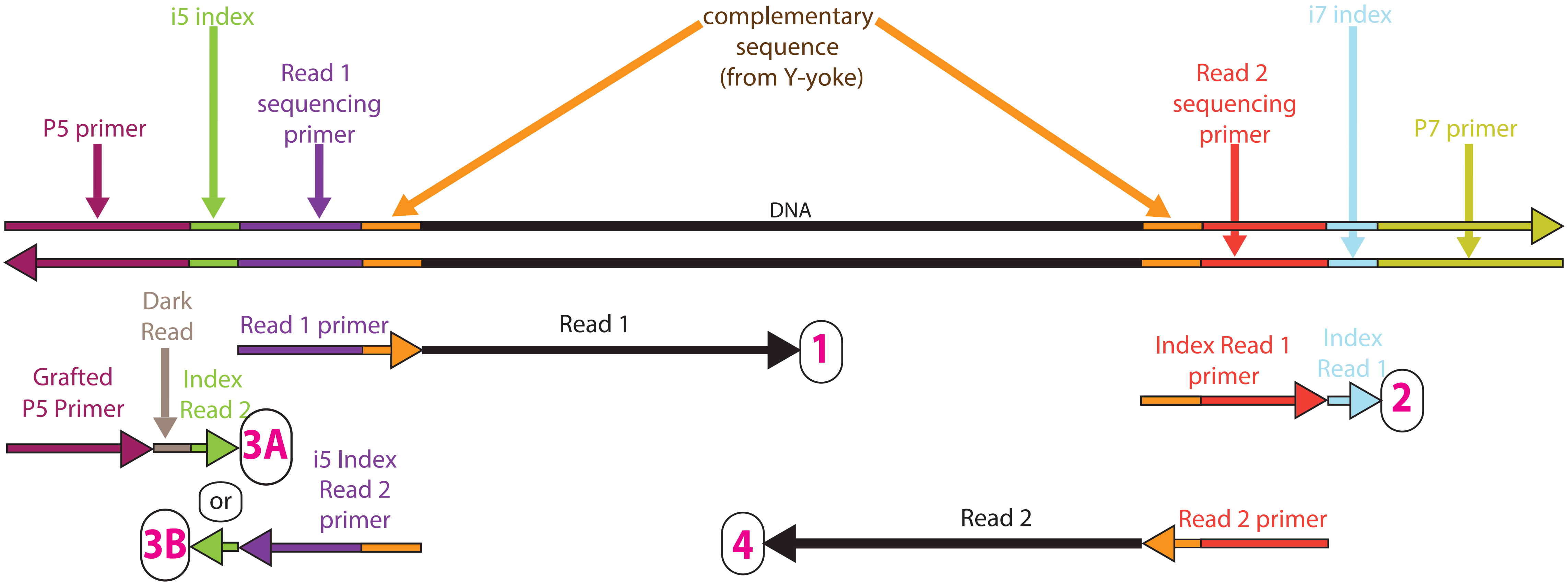

### Fig. S7

#### Sequencing Reads from Libraries Lacking an i5

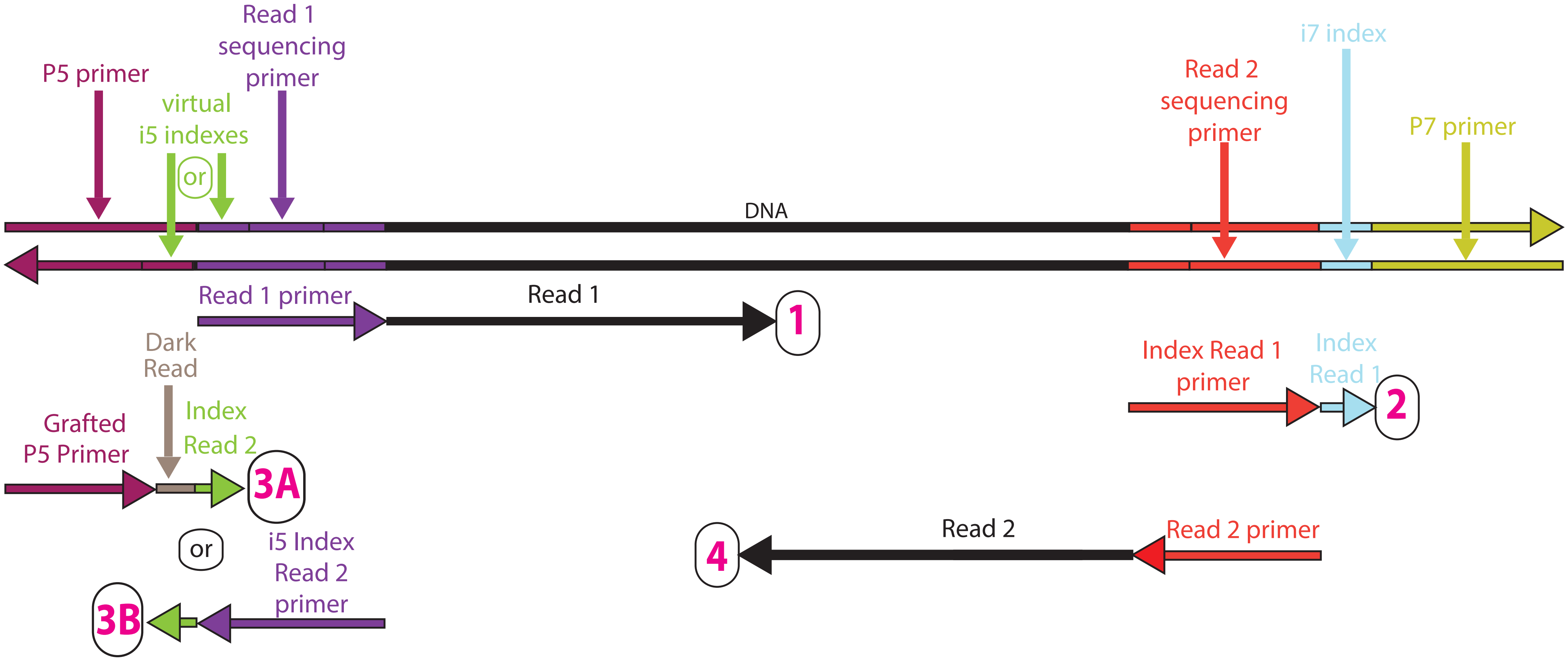

### Fig. S8

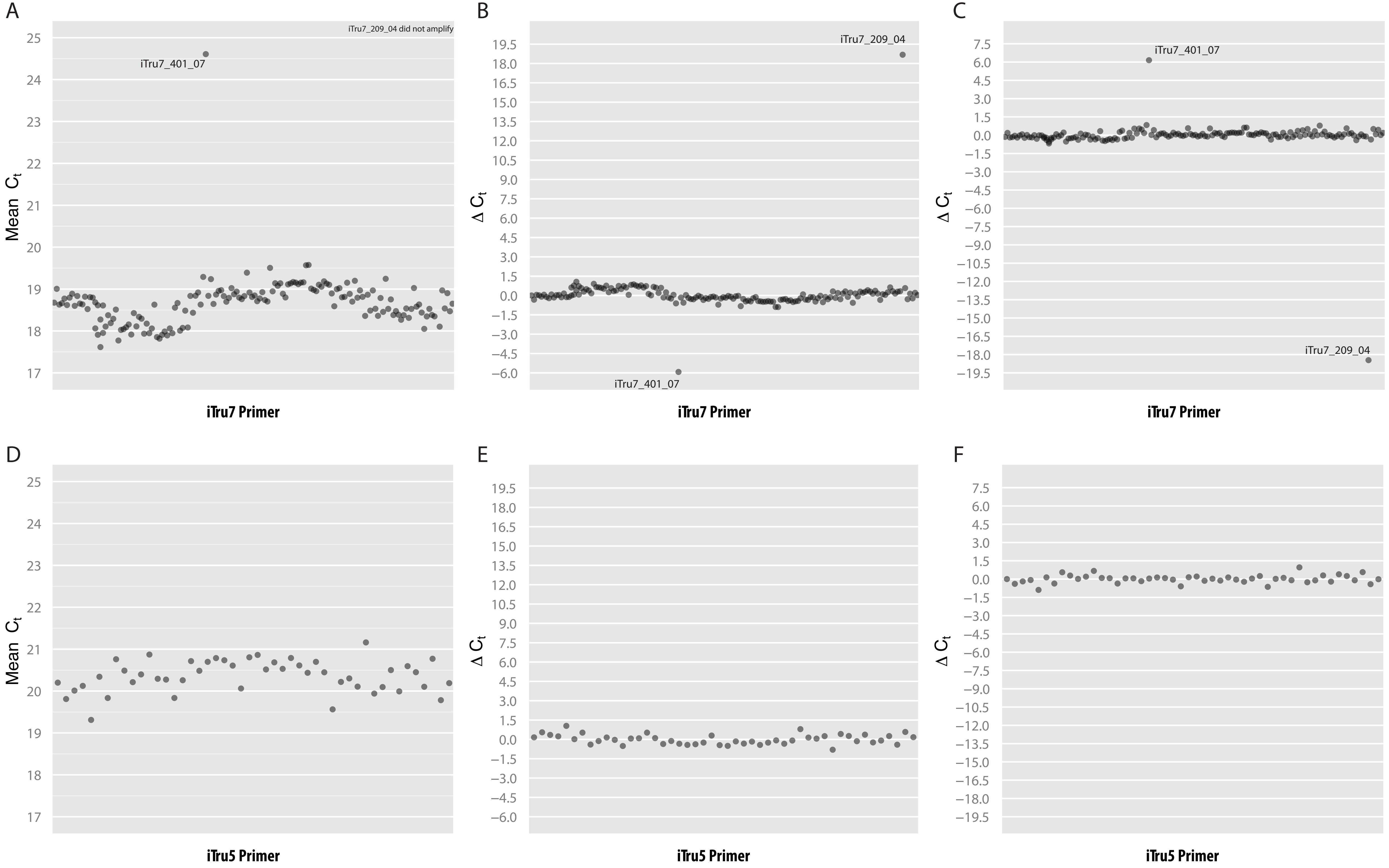

### Fig. S9

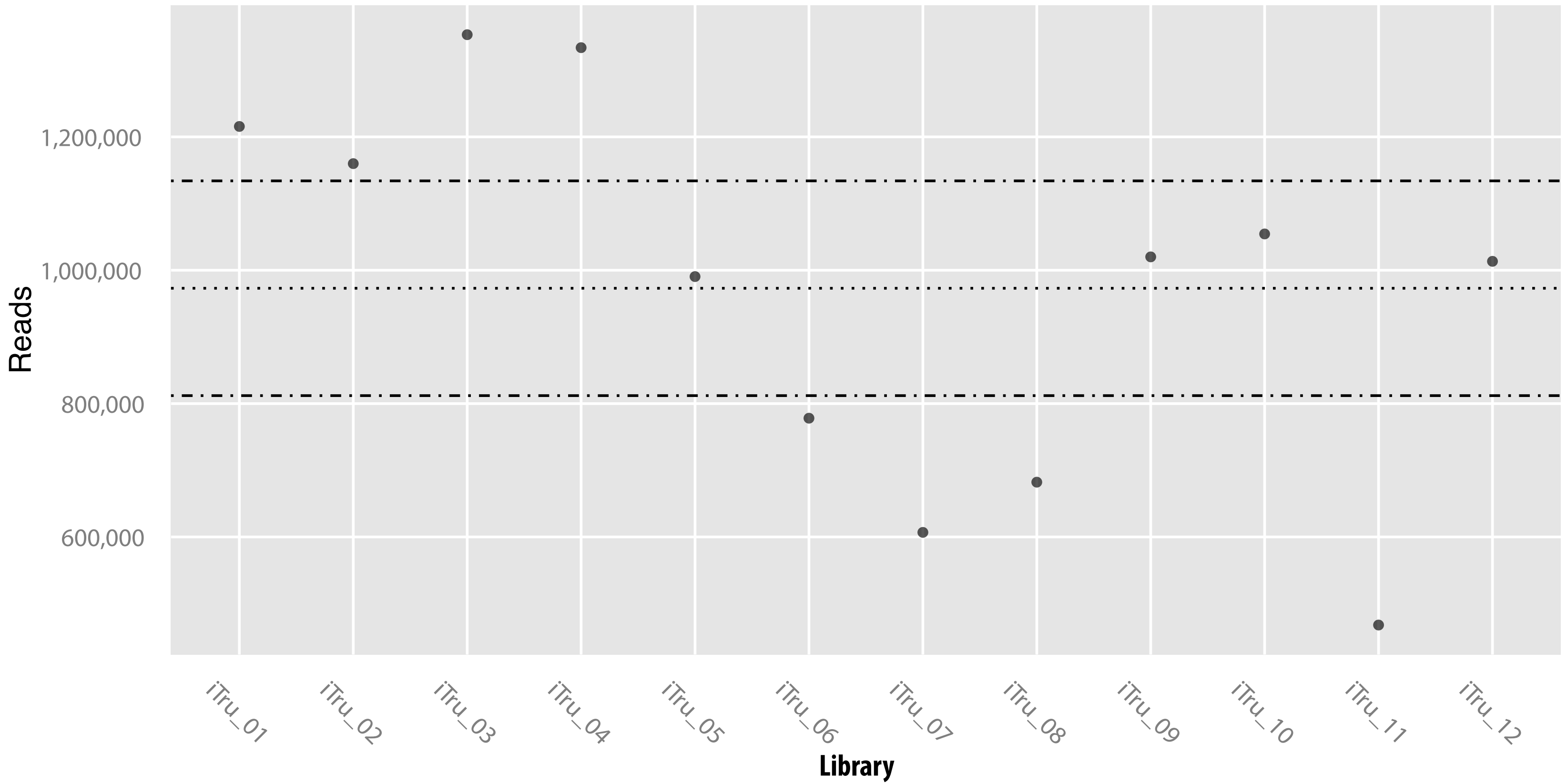

### Fig. S10

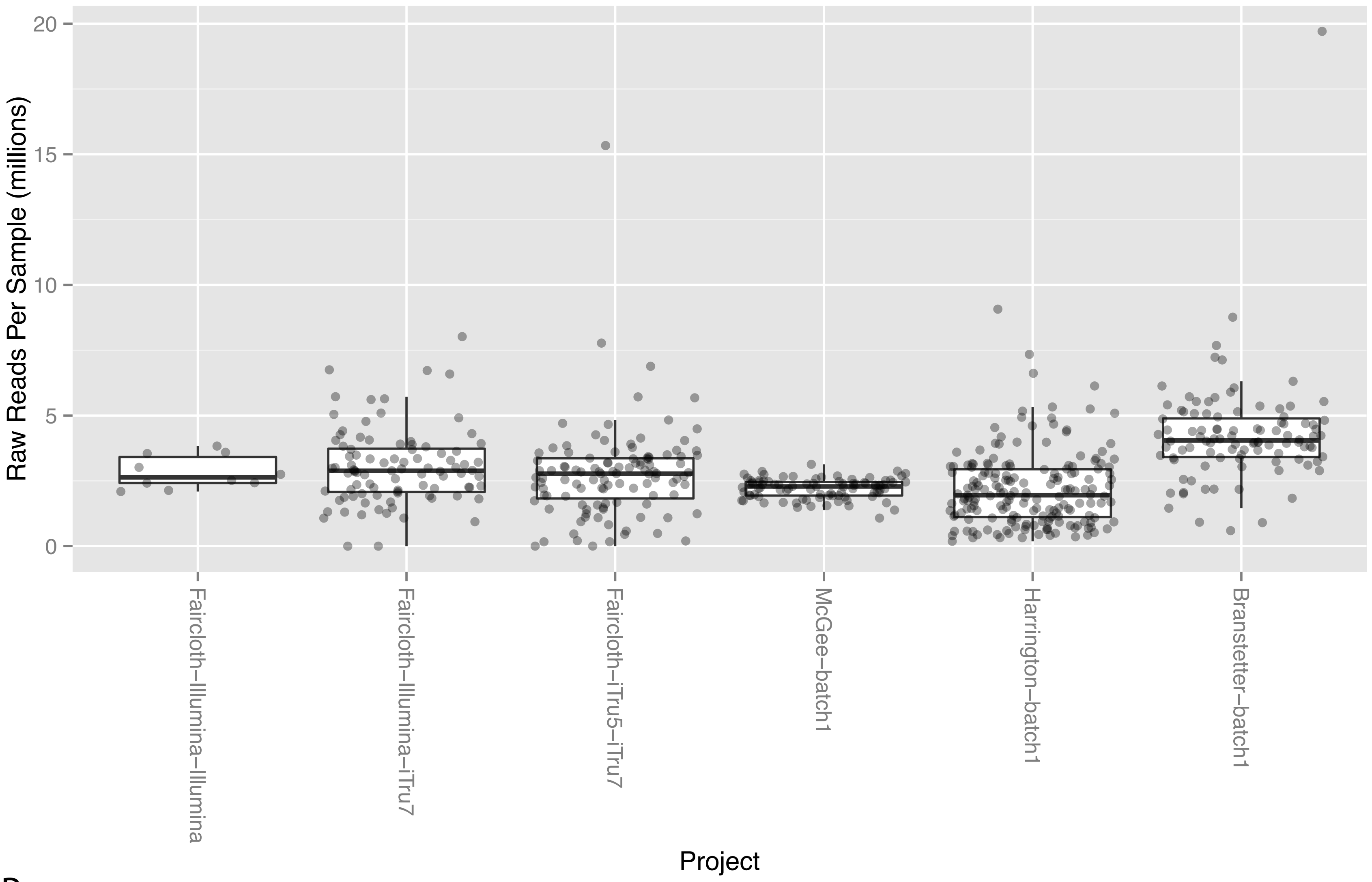

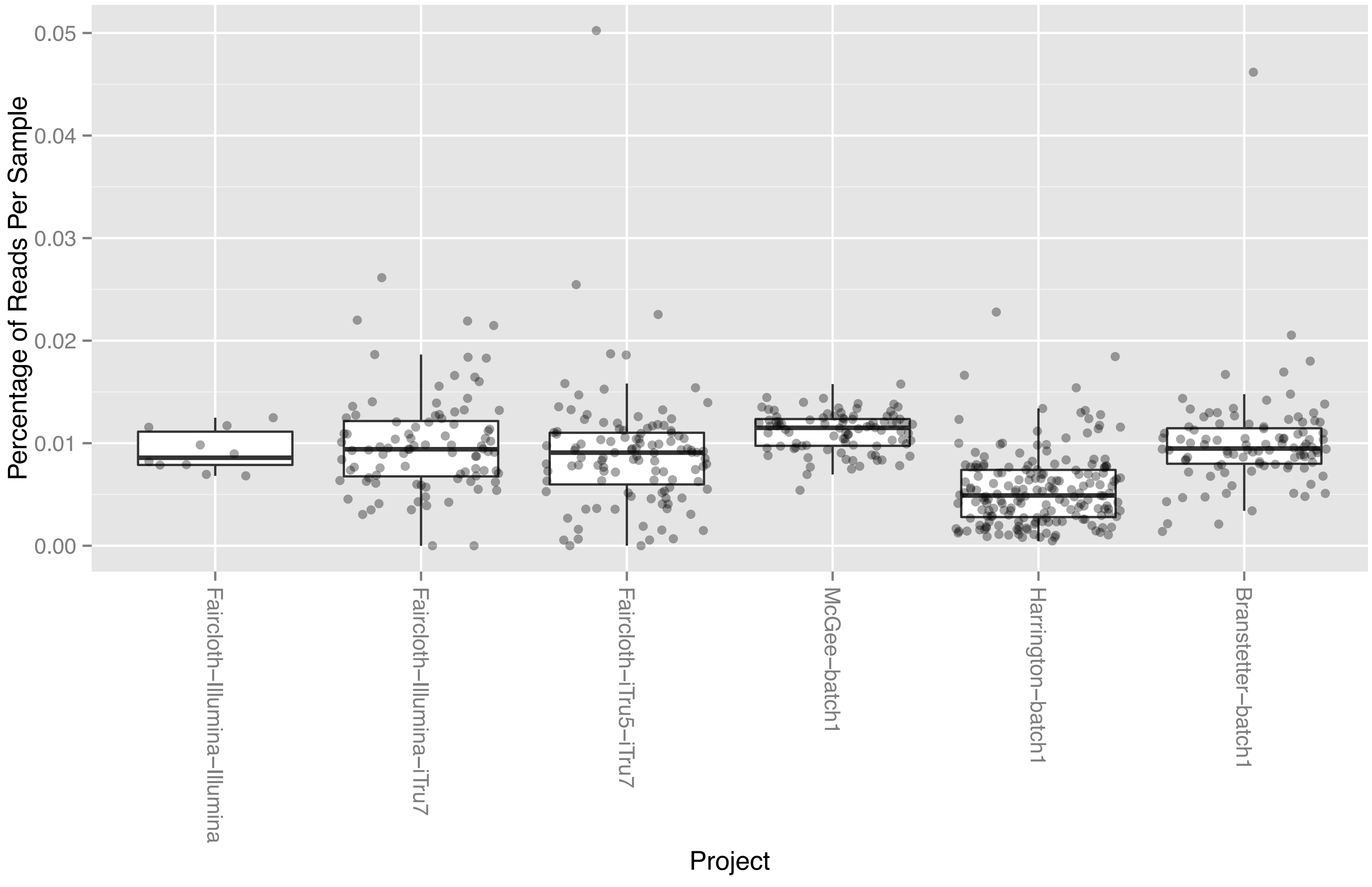

### Fig. S11

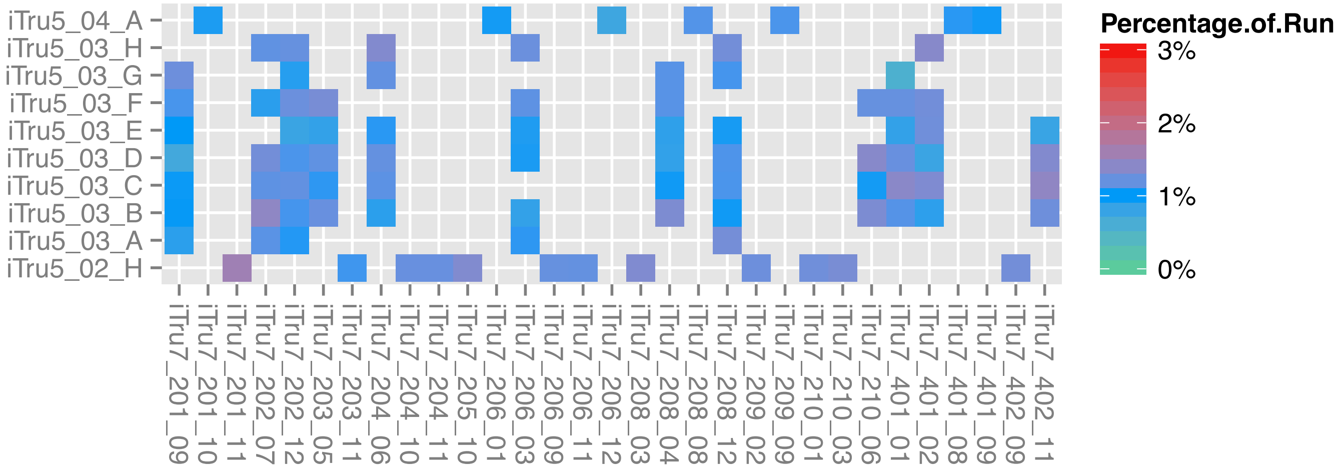

### Fig. S12

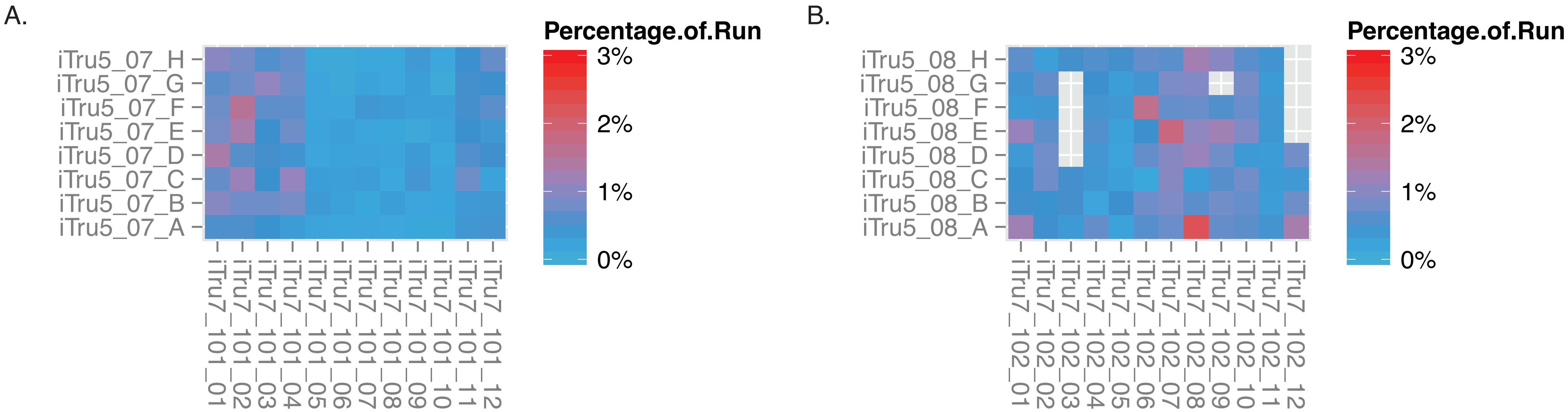

### Fig. S13

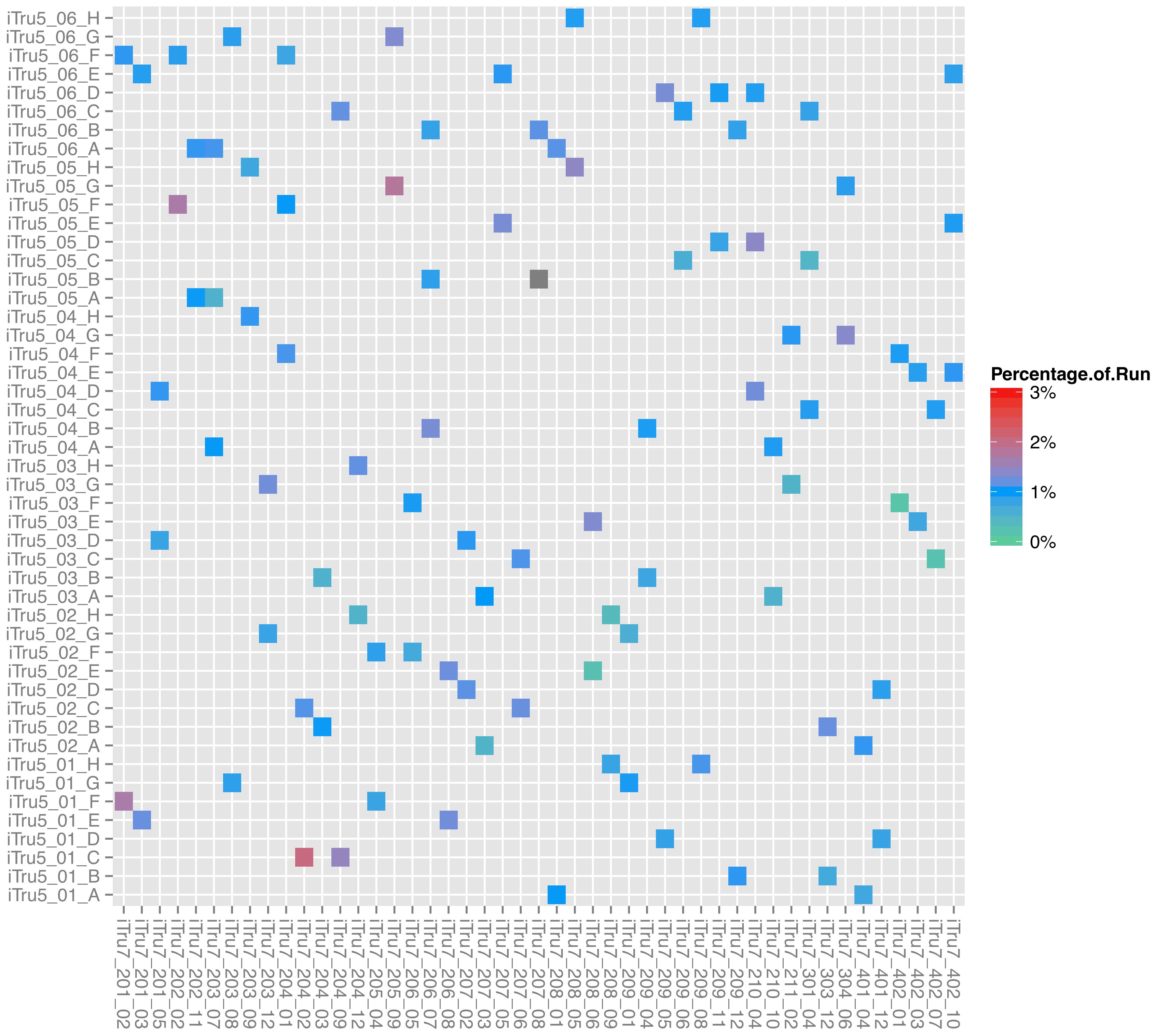

### Fig. S14

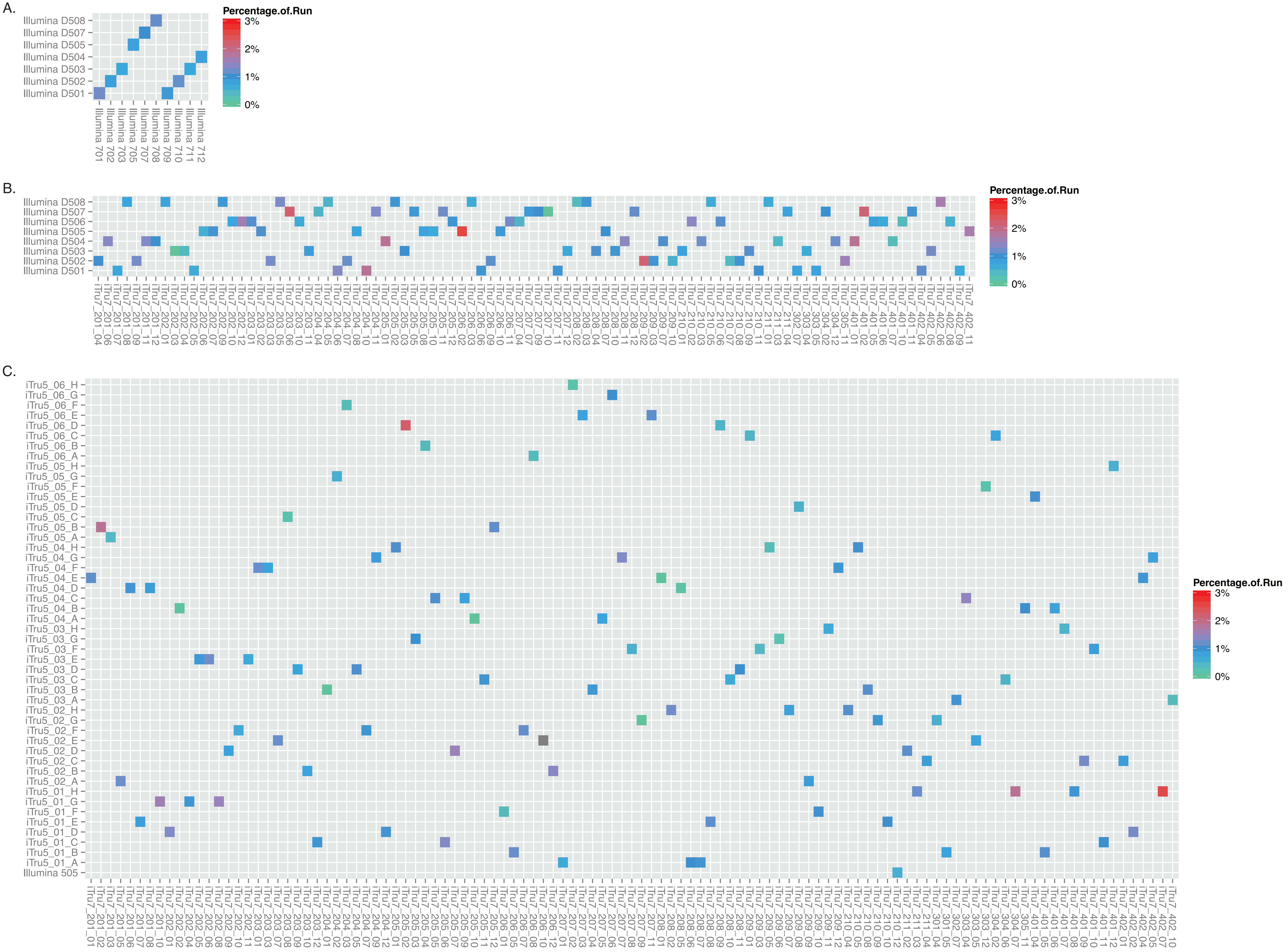

### Fig. S15

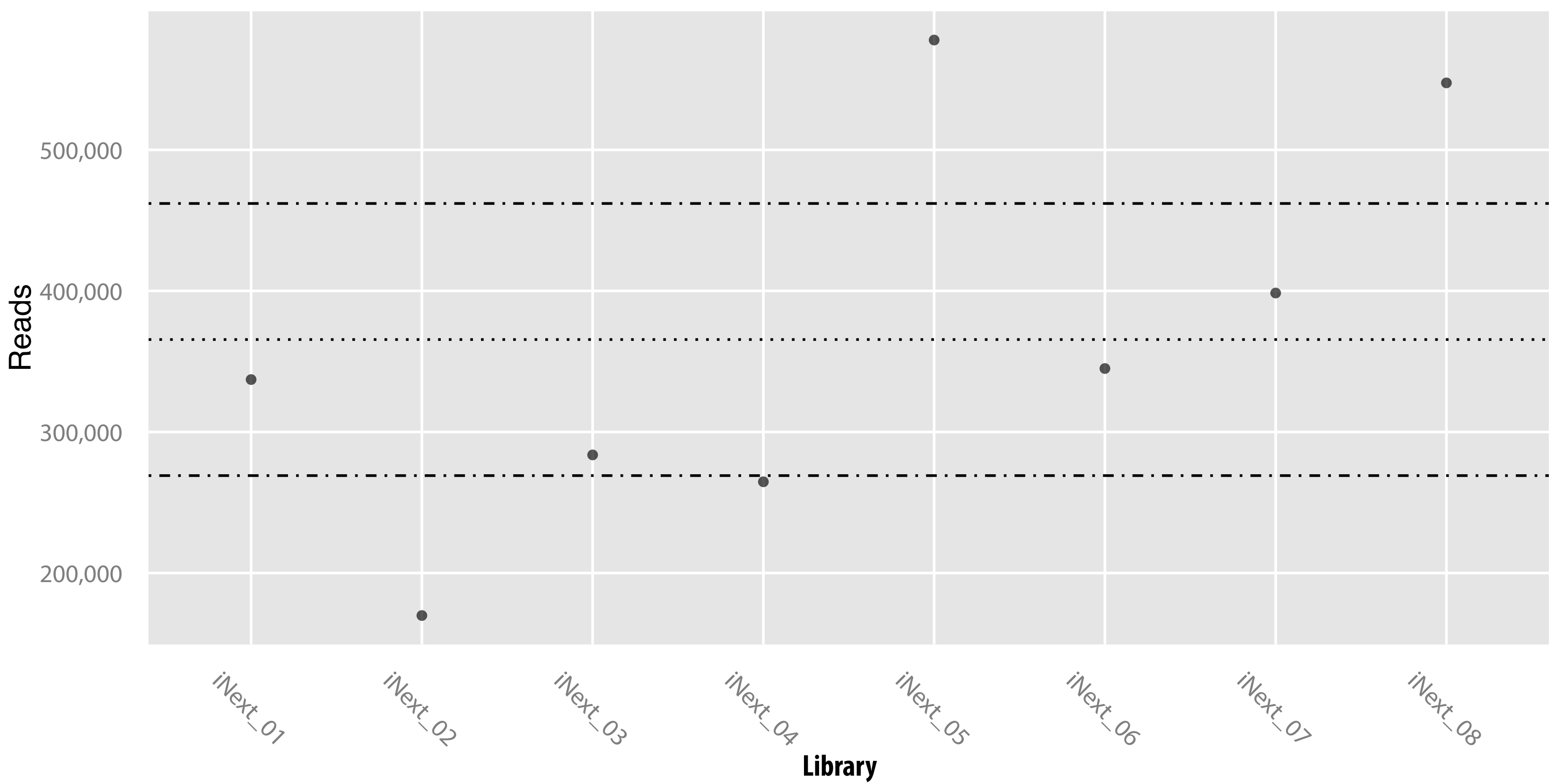
