## Supplementary material for "Adapterama I: Universal stubs and primers for 384 unique dual-indexed or 147,456 combinatorially-indexed Illumina libraries (iTru & iNext)": Fig. S5

| | D701 | D702 | D703 | D704 | D705 | D706 | D707 | D708 | D709 | D710 | D711 | D712 | TS–1 | TS–2 | TS–3 | TS–4 | TS–5 | TS–6 | TS–7 | TS–8 | TS–9 | TS–10 | TS–11 | TS–12 | TS–13 | TS–14 | TS–15 | TS–16 | TS–18 | TS–19 | TS–20 | TS–21 | TS–22 | TS–23 | TS–25 | TS–27 |
|---|---|---|---|---|---|---|---|---|---|---|---|---|---|---|---|---|---|---|---|---|---|---|---|---|---|---|---|---|---|---|---|---|---|---|---|---|
| D701 | – | – | – | – | – | – | – | – | – | – | – | – | – | – | – | – | – | – | – | – | – | – | – | – | – | – | – | – | – | – | – | – | – | – | – | – |
| D702 | 7 | – | – | – | – | – | – | – | – | – | – | – | – | – | – | – | – | – | – | – | – | – | – | – | – | – | – | – | – | – | – | – | – | – | – | – |
| D703 | 7 | 6 | – | – | – | – | – | – | – | – | – | – | – | – | – | – | – | – | – | – | – | – | – | – | – | – | – | – | – | – | – | – | – | – | – | – |
| D704 | 5 | 8 | 6 | – | – | – | – | – | – | – | – | – | – | – | – | – | – | – | – | – | – | – | – | – | – | – | – | – | – | – | – | – | – | – | – | – |
| D705 | 5 | 4 | 6 | 7 | – | – | – | – | – | – | – | – | – | – | – | – | – | – | – | – | – | – | – | – | – | – | – | – | – | – | – | – | – | – | – | – |
| D706 | 5 | 7 | 6 | 3 | 5 | – | – | – | – | – | – | – | – | – | – | – | – | – | – | – | – | – | – | – | – | – | – | – | – | – | – | – | – | – | – | – |
| D707 | 5 | 5 | 6 | 5 | 5 | 5 | – | – | – | – | – | – | – | – | – | – | – | – | – | – | – | – | – | – | – | – | – | – | – | – | – | – | – | – | – | – |
| D708 | 5 | 5 | 7 | 4 | 6 | 3 | 5 | – | – | – | – | – | – | – | – | – | – | – | – | – | – | – | – | – | – | – | – | – | – | – | – | – | – | – | – | – |
| D709 | 6 | 6 | 3 | 6 | 7 | 6 | 6 | 7 | – | – | – | – | – | – | – | – | – | – | – | – | – | – | – | – | – | – | – | – | – | – | – | – | – | – | – | – |
| D710 | 6 | 2 | 6 | 8 | 4 | 7 | 6 | 5 | 5 | – | – | – | – | – | – | – | – | – | – | – | – | – | – | – | – | – | – | – | – | – | – | – | – | – | – | – |
| D711 | 5 | 4 | 6 | 7 | 6 | 6 | 5 | 3 | 7 | 3 | – | – | – | – | – | – | – | – | – | – | – | – | – | – | – | – | – | – | – | – | – | – | – | – | – | – |
| D712 | 5 | 5 | 5 | 5 | 5 | 6 | 6 | 7 | 4 | 5 | 7 | – | – | – | – | – | – | – | – | – | – | – | – | – | – | – | – | – | – | – | – | – | – | – | – | – |
| TS–1 | 4 | 5 | 5 | 7 | 3 | 5 | 4 | 6 | 6 | 4 | 5 | 5 | – | – | – | – | – | – | – | – | – | – | – | – | – | – | – | – | – | – | – | – | – | – | – | – |
| TS–2 | 6 | 6 | 4 | 5 | 7 | 5 | 4 | 5 | 4 | 6 | 7 | 5 | 5 | – | – | – | – | – | – | – | – | – | – | – | – | – | – | – | – | – | – | – | – | – | – | – |
| TS–3 | 5 | 5 | 6 | 7 | 5 | 5 | 4 | 4 | 5 | 5 | 5 | 6 | 4 | 4 | – | – | – | – | – | – | – | – | – | – | – | – | – | – | – | – | – | – | – | – | – | – |
| TS–4 | 6 | 6 | 4 | 6 | 6 | 6 | 5 | 6 | 5 | 5 | 6 | 5 | 4 | 4 | 5 | – | – | – | – | – | – | – | – | – | – | – | – | – | – | – | – | – | – | – | – | – |
| TS–5 | 6 | 5 | 5 | 6 | 5 | 5 | 5 | 6 | 5 | 4 | 6 | 5 | 3 | 4 | 4 | 5 | – | – | – | – | – | – | – | – | – | – | – | – | – | – | – | – | – | – | – | – |
| TS–6 | 6 | 5 | 4 | 5 | 6 | 5 | 5 | 7 | 5 | 5 | 7 | 4 | 4 | 4 | 5 | 5 | 6 | – | – | – | – | – | – | – | – | – | – | – | – | – | – | – | – | – | – | – |
| TS–7 | 6 | 6 | 3 | 4 | 6 | 4 | 4 | 5 | 4 | 6 | 7 | 5 | 5 | 3 | 6 | 5 | 3 | 6 | – | – | – | – | – | – | – | – | – | – | – | – | – | – | – | – | – | – |
| TS–8 | 5 | 5 | 5 | 7 | 4 | 5 | 4 | 6 | 6 | 5 | 5 | 5 | 4 | 4 | 4 | 5 | 3 | 5 | 5 | – | – | – | – | – | – | – | – | – | – | – | – | – | – | – | – | – |
| TS–9 | 6 | 6 | 4 | 6 | 3 | 4 | 5 | 6 | 6 | 6 | 6 | 5 | 2 | 4 | 4 | 5 | 6 | 3 | 4 | 6 | – | – | – | – | – | – | – | – | – | – | – | – | – | – | – | – |
| TS–10 | 6 | 6 | 4 | 4 | 6 | 5 | 6 | 6 | 4 | 5 | 6 | 4 | 5 | 4 | 6 | 4 | 4 | 6 | 3 | 4 | 6 | – | – | – | – | – | – | – | – | – | – | – | – | – | – | – |
| TS–11 | 6 | 6 | 3 | 5 | 6 | 4 | 6 | 6 | 4 | 6 | 6 | 4 | 4 | 4 | 4 | 4 | 3 | 4 | 6 | 2 | 3 | 5 | – | – | – | – | – | – | – | – | – | – | – | – | – | – |
| TS–12 | 6 | 6 | 5 | 7 | 5 | 6 | 4 | 7 | 5 | 6 | 6 | 6 | 5 | 3 | 5 | 5 | 3 | 5 | 5 | 5 | 4 | 4 | 4 | – | – | – | – | – | – | – | – | – | – | – | – | – |
| TS–13 | 5 | 6 | 5 | 6 | 3 | 7 | 5 | 6 | 5 | 6 | 6 | 4 | 4 | 6 | 5 | 3 | 4 | 4 | 4 | 5 | 5 | 4 | 4 | 5 | – | – | – | – | – | – | – | – | – | – | – | – |
| TS–14 | 4 | 7 | 4 | 4 | 4 | 3 | 6 | 5 | 6 | 6 | 5 | 6 | 4 | 5 | 4 | 4 | 4 | 5 | 3 | 4 | 4 | 4 | 4 | 5 | 5 | – | – | – | – | – | – | – | – | – | – | – |
| TS–15 | 5 | 5 | 5 | 7 | 2 | 5 | 5 | 5 | 6 | 6 | 5 | 5 | 4 | 5 | 5 | 4 | 6 | 3 | 4 | 5 | 4 | 4 | 6 | 4 | 5 | 4 | – | – | – | – | – | – | – | – | – | – |
| TS–16 | 5 | 6 | 5 | 5 | 7 | 6 | 5 | 6 | 5 | 5 | 4 | 6 | 6 | 6 | 4 | 5 | 4 | 5 | 4 | 2 | 5 | 4 | 5 | 5 | 5 | 5 | 5 | – | – | – | – | – | – | – | – | – |
| TS–18 | 6 | 4 | 6 | 6 | 5 | 6 | 6 | 5 | 6 | 3 | 3 | 6 | 4 | 6 | 6 | 5 | 6 | 4 | 5 | 5 | 5 | 6 | 4 | 6 | 5 | 5 | 5 | 5 | – | – | – | – | – | – | – | – |
| TS–19 | 4 | 7 | 7 | 4 | 6 | 5 | 3 | 5 | 5 | 7 | 6 | 4 | 6 | 6 | 6 | 5 | 6 | 7 | 4 | 4 | 4 | 5 | 4 | 6 | 5 | 3 | 5 | 6 | 4 | – | – | – | – | – | – | – |
| TS–20 | 6 | 7 | 5 | 6 | 7 | 5 | 5 | 7 | 5 | 6 | 6 | 7 | 5 | 6 | 5 | 5 | 5 | 6 | 5 | 5 | 4 | 6 | 6 | 5 | 3 | 4 | 5 | 7 | 5 | 6 | – | – | – | – | – | – |
| TS–21 | 5 | 5 | 6 | 6 | 4 | 4 | 6 | 6 | 6 | 5 | 4 | 7 | 5 | 6 | 7 | 6 | 6 | 7 | 5 | 6 | 7 | 7 | 6 | 6 | 5 | 4 | 5 | 6 | 6 | 6 | 5 | – | – | – | – | – |
| TS–22 | 5 | 5 | 4 | 7 | 5 | 5 | 5 | 6 | 5 | 5 | 6 | 5 | 5 | 3 | 4 | 5 | 6 | 5 | 6 | 6 | 4 | 6 | 4 | 6 | 4 | 5 | 5 | 5 | 6 | 5 | 5 | 4 | – | – | – | – |
| TS–23 | 7 | 6 | 6 | 5 | 5 | 4 | 5 | 5 | 6 | 6 | 7 | 5 | 5 | 3 | 6 | 5 | 7 | 5 | 6 | 6 | 5 | 7 | 5 | 5 | 4 | 6 | 5 | 5 | 5 | 7 | 6 | 4 | 6 | – | – | – |
| TS–25 | 5 | 5 | 4 | 6 | 4 | 6 | 4 | 6 | 4 | 5 | 6 | 3 | 4 | 4 | 4 | 4 | 6 | 6 | 5 | 6 | 6 | 4 | 5 | 5 | 6 | 4 | 5 | 5 | 5 | 5 | 4 | 7 | 6 | 4 | – | – |
| TS–27 | 3 | 7 | 5 | 6 | 4 | 5 | 6 | 6 | 6 | 6 | 6 | 6 | 4 | 6 | 5 | 6 | 7 | 7 | 7 | 5 | 4 | 6 | 6 | 6 | 6 | 4 | 3 | 6 | 4 | 6 | 5 | 5 | 5 | 6 | 4 | – |
