## Supplementary material for "Adapterama I: Universal stubs and primers for 384 unique dual-indexed or 147,456 combinatorially-indexed Illumina libraries (iTru & iNext)": File S1

Supplemental File 1: iNext Details

Adapterama I: Universal stubs and primers for 384 unique dual-indexed or 147,456 combinatorically-indexed Illumina libraries (iTru & iNext)

**Introduction**

Similar to our modifications of the Illumina TruSeq system, we modularized the Illumina Nextera system by dividing by dividing the adapter components into two parts: a universal Y-yoke adapter stub that comprises most of the Read 1 and 2 primer binding sites (Table 3; Supplemental File 15) and a set of amplification primers (iNext5, iNext7; described below), parts of which are complementary to the Y-yoke stub and which also contain custom sequence tag(s) for sample indexing (Figs. S2; Supplemental File 15). It is **important to note** that the iNext Y-yoke adapter has a single 5’ guanosine (G) overhang and thus requires library preparations that produce insert DNA with single 3’ cytidine (C) overhangs to ensure proper base-pairing and ligation.

Similar to previous custom Nextera primers that we designed to extend the Epicentre Nextera system (Crawford et al., 2012), we designed a large set of indexed amplification primers (iNext5, iNext7) that contain a subset of our custom 8 nt sequence tags from Faircloth & Glenn (2012; File S15). All tags used are compatible with 8 nt tags in the original Illumina Nextera primers and maintain an edit distance ≥3 to allow error correction. Similar to the iTru primers, we grouped the iNext primers groups into clearly identifiable numbered sets with eight primers in iNext5 sets and 12 primers iNext7 sets. Each set of primers has balanced index base composition (Illumina, 2016).

We ordered the components of our iNext Y-yoke adapter stubs and iNext primers from Integrated DNA Technologies (IDT, Coralville, IA, USA). We modified the adapter stub sequence by phosphorylating the 5’ end of iNext_R2_stub_RCp, and we modified each of the iNext primer sequences by adding a phosphorothioate bond before the 3’ nucleotide of each sequence to inhibit degradation due to exonuclease activity (Skerra, 1992).

In contrast to Illumina Nextera kits, these iNext Y-yoke adapters and iNext primers allow us to produce Nextera-style libraries (i.e., iNext libraries) at low-cost per sample when reasonably large quantities of DNA (e.g., > 100ng) are available, when specific insert-size distributions are necessary, or when all available input DNA is highly fragmented. We continue to use Illumina Nextera or Nextera XT kits when limited quantities of intact DNA are available and when broader insert-size distributions are acceptable. The primary limitation of iNext libraries relative to iTru libraries is the widespread use of TruSeq libraries and the fact that Illumina advises against combining Nextera and TruSeq libraries in the same sequencing lane (Illumina, 2012).

It is important to note that because the Nextera and TruSeqHT indexes are not the same, the subset of tags incorporated into iNext and iTru are not identical. Further, there is no correspondence of tags between iNext and iTru (e.g., iNext5_01_A is not the same tag as iTru5_01_A). See additional information on this topic in the Discussion of the main manuscript.

**Materials & Methods**

***Validation of iNext Primers by Quantitative PCR (qPCR)***

Similar to our validation of the iTru primers, we formed qPCR reactions with some of our iNext primers to determine if any of the indexed primers were biasing amplification. We selected a subset of iNext5 and iNext7 primers for qPCR validation (File S15, 2012_iNext5_information, 2012_iNext7_information), and we setup qPCR reactions that were identical to those we used to test the iTru primers. We tested all forward (iNext5) primers with Illumina N701 as the reverse primer and we tested all reverse primers (iNext7) with Illumina N501 as the forward primer. As with the iTru primer tests, we evaluated the relative performance of all iNext5 and iNext7 primers by subtracting the C_T_ of the iNext5 or iNext7 primer being tested from the average C_T_ of all iNext5 or iNext7 primers. Second, we evaluated the performance of all iNext5 and iNext7 primers by subtracting the baseline reference C_T_ of iNext5_01_A from the C_T_ of the iNext5 primer being tested and by subtracting the baseline reference C_T_ of iNext7_01_01 from the C_T_ of the iNext7 primer being tested. We expected that unbiased primers would not deviate from the average and/or baseline performance by more than 1.5 PCR cycles.

***Implementation: DNA source***

To test the performance of both our iNext Y-yoke adapters and the iNext system in real-world library preparation scenarios, we prepared libraries from DNA of various types and quality. As a source of control DNA, we used *Escherichia coli* k-12 strain MG1655 (*E. coli* hereafter; Roche, Basel, Switzerland). To examine how our iNext system performed with DNA of varying quality and complexity, we also prepared iNext libraries from DNA that we obtained from two *Blastomyces dermatitidis* clinical strains (Marshfield Labs, Marshfield, WI, USA). Both strains were isolated from human patients in 2000 using extraction procedures described in Meece *et al.* (2010). *Blastomyces dermatitidis* is a fungal pathogen of humans with an incompletely sequenced genome (the genome assembly contains thousands of scaffolds with limited linkage information among the scaffolds). We refer to these *Blastomyces dermatitidis* DNA extracts as the “fungal” samples.

***Implementation: iNext library construction***

We prepared eight *E. coli* and two fungal DNA sample using a similar approach to what we used in the main manuscript text, except that we replaced the 10X A-Tailing buffer (KB8211) of the Kapa library kit with NEB buffer 2 (New England Biolabs, Beverly, MA, USA) supplemented with 1 mM dCTPs (Sigma, St. Louis, MO, USA), and we used iNext universal adapter stubs during the ligation. We created full iNext library constructs (Fig. S4) by performing a subsequent limited-cycle PCR reaction using the iNext primers (Table 3; Supplemental File 15). Step-by-step methods are outlined in Supplemental File 16.

***Implementation: Nextera XT library library construction***

Illumina Nextera XT kits were used to construct libraries for both fungal samples as well as the set of eukaryotic species given in Table 4 of the main text. For library construction, we adhered to the manufacturer’s instructions with the following exceptions: 1) reactions were completed at half volume, 2) DNA was at 2 ng/µL (instead of 1 ng/µL) to increase the average insert size, 3) iNext primers were used, 4) an additional 4 cycles of PCR were needed, and 5) following PCR, the libraries were checked on an agarose gel, cleaned, quantified, and normalized using similar techniques as were used for iTru (i.e., we did not use the Illumina reagents for purification and normalization of the Nextera XT libraries).

***Sequencing***

iNext libraries from the E. coli were pooled with iTru E. coli libraries, as well as unrelated samples prepared using Illumina TruSeq library preparation kits, and sequenced on a MiSeq at the University of Georgia using v2 500 cycle run to produce PE250 reads. Fungal libraries (iNext and Nextera XT) were pooled and sequenced on a MiSeq at the Marshfield Clinic using v2 500 cycle run to produce PE250 reads.

***Sequence Analysis***

We mapped *E. coli* reads back to NC_000913 using the Geneious mapper (fastest setting, single iteration). Fungal reads were mapped to ATCC26199 and SLH14081 using the Geneious mapper (fastest setting, single iteration) in Geneious version R6.1.7.

**Results**

***Quantitative PCR***

All but one of the iNext primers (47/48 iNext7’s and 48/48 iNext5’s) had average C_T_ values within 1.5 cycles of the average C_T_.

***iNext libraries***

iNext *E. coli* library preparations produced the expected smear pattern on an agarose gel , but required 2 additional cycles of PCR to yield similar concentration as iTru *E. coli* libraries.

Roughly similar numbers of reads were obtained among the iNext *E. coli* libraries (averaging 365,537 reads per sample; supplemental file 13). All of the libraries produced high quality reads with mapping rates of 95-96%, averaging 303,997 reads mapped per sample. All of the samples covered >99% of the known *E. coli* genome sequence except the single sample with the lowest number of reads (169,868) which only covered 98.4% of the known genome sequence. Although the number of reads among iNext replicates were similar, as were the iTru replicates, fewer reads were obtained on average for iNext than iTru (365,537 vs. 973,008).

***Fungal iNext & NexteraXT***

A limited number of reads were obtained from the NexteraXT libraries, with high numbers of PCR duplicates (data not shown). Clearly, using half-sized reactions of NexteraXT for fungal genomic libraries was too far from the recommended use of the product and thus, NexteraXT libraries were not analyzed further. For the two strains of fungi (samples 26 and 33), a large number of reads were obtained from the MiSeq run (9,556,598 and 16,750,472 for samples 26 and 33, respectively; supplemental file 14)..Most of the reads were ≥200 bases. The vast majority of sites mapped with high quality (>98% with ≥Q30 for all combinations) and most of the sites were identical between the samples and reference sequences (84.4% to 96.5%). However, the percentage of the reference genomes covered by mapped reads varied from 77.7% to 97.8%, which reflects the divergence between the samples and the references.

**Discussion**

iNext library preparations produced libraries that easily passed QC metrics. These libraries were pooled with NexteraXT as well as iTru libraries and successfully sequenced on the MiSeq. iNext libraries produced the expected number of reads which evenly cover the genome. As should have been the case due to the much higher amount of input DNA, iNext libraries were more diverse than libraries produced using NexteraXT. Thus, the iNext stubs presented here will give researchers an additional option for producing high quality libraries at low cost.

There are, however, a few limitations that researchers should keep in mind prior to choosing iNext. First, much more DNA is required for iNext than for NexteraXT or Nextera. Second, iNext library preparation requires much more time than a Nextera or NexteraXT library prepation. Third, about 2 more cycles of PCR are required to produce a yield similar to iTru libraries, and even with these 2 extra cycles, only about one-third as many reads were obtained from the iNext libraries. Thus, it appears that the number of molecules that are successfully ligated is lower for iNext than iTru, which will lead to libraries of slightly lower diversity. Our further efforts, therefore, focused on iTru (see main text).

***Pooling iNext with iTru libraries***

Illumina strongly advises against combining Nextera and TruSeq libraries within MiSeq runs and HiSeq lanes, but does support Nextera and TruSeq libraries being sequenced in separate lanes of HiSeq flow cells (Illumina, 2012; most recently accessed 14 June 2016). Running both types of libraries on the MiSeq seemed feasible to us because Illumina provides both TruSeq and Nextera sequencing primers within the same wells of the MiSeq kits, thus both types of primers are within all MiSeq runs that use the standard Illumina primers. The HiSeq is somewhat different in that primers are always user-supplied. Thus, it is not clear whether pooling both library types within a HiSeq lane would successfully produce sequences. Because of the high cost of conducting such experiments, we are not likely to undertake pooling experiments on the HiSeq.

It is important to note that even with increased PCR cycles to boost iNext library concentrations to be similar to iTru, only about one-third as many reads were obtained from the iNext libraries. The reduced number of reads may be due to incorrect quantification of the iNext pool and/or bias in cluster generation and/or sequencing efficiency of the sequencing primers. It seems most likely that the difference is due to quantification error, but we cannot rule out the other possibilities.

Due to our success in combining iNext and iTru libraries within the MiSeq run reported here, we are now routinely combining iNext-style and iTru-style libraries within MiSeq runs. We have combined iTru and iNext libraries within several runs on the MiSeq and have obtained the expected/desired results in all cases (data only shown). It is important to note, however, that in all of these runs, the iNext-style libraries are a minor component of the run (≤10%). We also note that these pooled runs are becoming much less frequent since the release Hyper and Hyper Plus kits from Kapa Biosciences, which opens up the use of iTru primers for very small amounts of starting DNA.

We do not guarantee that this will work for everyone under all possible scenarios, but we do encourage Illumina to undertake new work to determine under what conditions combining TruSeq-style and Nextera-style libraries can be done successfully. If Illumina is unable to do these tests internally, we would encourage them to provide sufficient reagents to research community members to test this important aspect of their technology.

A primary consideration when pooling libraries is that the tags on all samples should be unique. The iNext and iTru designs draw from the same pool of tags, but because the tags used in the Illumina primers for Nextera and TruSeqHT are not the same, the iNext and iTru primer tag sets are not identical (i.e., they both draw from the 531 8nt tags of Faircloth and Glenn [2012], given in supplemental file 4; but because Nextera and TruSeqHT indexes are different, the sets of non-conflicting 8 nt tags are different). Additionally, as we note in the introduction, the iNext and iTru primer sets do not have correspondingly equivalent tags (e.g., the tag used in iNext_i5_01_A is NOT equivalent to iTru_i5_01_A). Unique tag identifiers (from Faircloth & Glenn [2012]) are given in supplemental files 4 and 15, which must be consulted to match up tags between iNext and iTru.

**Overview:** Here, we are MOSTLY following the methods for Kapa Hyper Prep library preparation kits. But, in addition to using half volumes and Speedbeads (as we do for the iTru library preps), we also have to make a custom buffer to add a single C (from dCTP) after end repair, rather than the usual A (from dATP). So, Step #6 is modified from the normal library preps, and we use the iNext Y-yoke adapter stub, otherwise it is the same as the iTru library prep.

**Protocol:**

1. Normalize DNA at ~23ng/µL
2. Biorupter fragmentation: 3 sets of 5 cycles (15cycles) on Medium setting.
3. Insert 25µL of sheared DNA in 10µL of End Repair Master Mix.
4. Incubate in Thermocycler at 20**°**C for 30 minutes.
5. Add 100µL of speedbeads to 35µL volume, mix and let sit for 5-15 minutes.
   - Magnet capture & EtOH wash twice with 80µL of 80% EtOH.
6. To dry beads add 25µL of C-tailing master mix [1x NEB Buffer 2 (10 mM Tris HCl pH7.9, 10 mM MgCl_2_, 50 mM NaCl, 1 mM DTT), 0.2 mM dCTP, 5 units Klenow Fragment (3’→5’ exo-; NEB M0212)].
7. Incubate in Thermocycler at 37**°**C for 30 minutes.
8. Add 45µL of PEG/NaCl to 25µL volume, mix and let sit for 5-15 minutes.
   - Magnet capture & EtOH wash twice with 80µL of 80% EtOH.
9. To dry beads add 22.5µL of Ligation master mix and 2.5µL of 5uM iNext Y-yoke adapters.
10. Incubate in Thermocycler at 20**°**C for 15 minutes.
11. Add 25µL of PEG/NaCl to 25µL volume, mix and let sit for 5-15 minutes.
    - Magnet capture & EtOH wash twice with 80µL of 80% EtOH.
12. To dry beads add 100µL of TLE (or IDTE; 10 mM Tris pH8, 0.1 mM EDTA).
13. Add 55µL of PEG/NaCl to 100µL volume, mix and let sit for 5-15 minutes.
14. Magnet capture transfer clear supernatant (~155µL) to new tube and discard the old speedbeads.
15. Add 25µL of speedbeads to 155µL volume of transferred supernatant, mix and let sit for 5-15 minutes.
    - Magnet capture & EtOH wash twice with 80µL of 80% EtOH
16. To dry beads add 25µL of TLE (IDTE). This is the Pre-PCR Library Prep.
17. PCR: To a new tube add 12.5µL of KAPA Hotstart Ready Mix, 1.25µL of i5 primer, 1.25µL of i7 primer, and 10µL of the Pre-PCR Library Prep. with speedbeads, mix and place in thermocycler for 8-12 cycles.
18. Run 5µL on a gel.
19. For samples that need it, do more cycles.
20. Cleanup: add 25µL of speedbeads to 20µL volume.
21. Quantify with Qubit.
