## Supplementary material for "Adapterama I: Universal stubs and primers for 384 unique dual-indexed or 147,456 combinatorially-indexed Illumina libraries (iTru & iNext)": File S6

When you receive the primers, there are 1.25 nmol of dried primer in each well. You will need to reconstitute them to the appropriate volume (125 µL -> 10 µM).

**Acceptable liquids to use for reconstitution are:**

1) 10 mM Tris pH 8 (or pH 7.5), or

2) TLE (homemade or IDTE from IDT = 10 mM Tris pH 8 & 0.1 mM EDTA)

**Protocol:**

1. Centrifuge the plates to get all the primer to the bottom of the wells.
2. To limit contamination, peel back the foil cover from the plate one row or column (depending on loading scheme) at a time to reconstitute.
3. Add 125 µL of your chosen liquid (e.g., TLE) to each well. Then, using a multichannel pipette:
   - Skloosh (i.e., pipette up & down) several times & use the pipet tip to help scrape the bottom of the well to dislodge any of the primer that is stuck.
   - Skloosh several more times.
   - Wait a few minutes and then skloosh several more times.
   - Let the primers sit in the liquid at room temperature for at least 5 minutes.
4. Once the primers have been fully dissolved, transfer the liquid to new strip tubes.
   - It is wise to use tubes of different color for each set. Be certain to maintain left/right orientation of the strips.
   - It is wise to aliquot each row/column into multiple strips (i.e., so you have <125 µL per primer in each tube). You want these primers to undergo freezing & thawing as few times as possible, so small aliquots are best (in practice 25 - 62.5 µL aliquots are reasonable).
5. Each primer is now at 10 µM and needs to be diluted down to 5 µM (25 µL TLE and 25 µL 10 µM primer for a 50 µL aliquot). Now they are ready to use.
   - It is wise to Nanodrop or otherwise quantify your primers. These primer aliquots are quite small, so they are easy to miss. If you miss the tiny dried amount in a well, your primers aren’t at 5 µM & your library PCRs won’t work!
   - Use 2.5 µL for each 50 µL reaction; you started with enough for 100 reactions.
6. Store the primers at -20°C
7. Take primers out to thaw shortly before use and skloosh well before using!
