## Supplementary material for "Adapterama I: Universal stubs and primers for 384 unique dual-indexed or 147,456 combinatorially-indexed Illumina libraries (iTru & iNext)": File S7

When you receive the oligos that you will use to make adapters (e.g., from IDT), if you get the full synthesis, there will be varying amounts in each tube or well of a plate. You need to reconstitute each oligo with the appropriate volume of liquid to achieve the desired concentration (i.e., 100 µM), then mix the two oligos that make the adapter (e.g., 50 µL each -> 100 µL at 50 µM), anneal them, then dilute again (to 5 µM) & aliquot.

To prepare the salty TLE for reconstitution & annealing (10 mM Tris pH 8, 0.1 mM EDTA, 100 mM NaCl) add the following to a 50 mL conical:

40 mL dH_2_O

500 µL 1M Tris pH 7.5 to 8

20 µL 0.5M EDTA pH 8

1 mL of 5 M NaCl

Fill with distilled water to 50 mL mark.

**Protocol:**

1. Centrifuge the plates or tubes to get all the primer to the bottom of the wells/tubes.
2. To limit contamination, peel back the foil cover from the plate one row or column (depending on loading scheme) at a time to reconstitute or deal with tubes individually. Use barrier (filtered) pipette tips for each step.
3. Add salty TLE at 10 times the number of nmol of oligo (e.g., if you have 79.2 nmol; add 792 µL of salty TLE).
   a. Use the pipette tip to help scrape the bottom of the tube/well to dislodge any of the oligo that is stuck.
   b. Skloosh (pipette up & down) several times until sufficiently mixed.
   c. Wait a few minutes.
   d. Skloosh several more times until sufficiently mixed.
   e. Let the oligos sit in the liquid at room temperature for at least 5 minutes.
4. Mix equal volumes of each adapter in the pair (50 µL each oligo -> 50 µM adapter) in strip tubes.

5. Anneal the adapters together: by placing the strip tube in a thermal cycler to denature (95 °C for 1 min.) & cool slowly (e.g., 0.1 °C per sec.) to room temp.
6. Dilute aliquots of the annealed adapters into new strip tubes by adding 10 µL of annealed adapters to 90 µL of salty TLE (final concentration = 5 µM; adjust this aliquot volume to be what you will typically use up while processing a single set of samples).
7. Store the strip tubes of adapters at -20 °C.
8. Take adapters out to thaw shortly before use, removing only the number of tubes needed (when appropriate, cut off individual tubes).
9. Once thawed, skloosh well before using!
10. Adapters can be refrozen but lose effectiveness with multiple freeze-thaw cycles. So, it is generally best to minimize the number of freeze-thaw cycles by making small aliquots that get used up quickly.
