## Supplementary material for "Adapterama I: Universal stubs and primers for 384 unique dual-indexed or 147,456 combinatorially-indexed Illumina libraries (iTru & iNext)": File S8

The goal here is to create a substitute for AMPure XP that is of equal effectiveness in comparison to the commercial product but far more cost-effective ($19/mL versus $0.46/mL).

**Suggested Reading**

Rohland N, Reich D. **Cost-effective, high-throughput DNA sequencing libraries for multiplexed target capture.** Genome Research 22: 939-946.

DeAngelis MM, Wang DG, Hawkins TL: **Solid-phase reversible immobilization for the isolation of PCR products**. *Nucleic Acids Res* 1995, **23**:4742–4743.

Fisher S, *et al.*: **A scalable, fully automated process for construction of sequence-ready human exome targeted capture libraries.** *Genome Biol* 2011, **12**:R1.

Lundin S, Stranneheim H, Pettersson E, Klevebring D, Lundeberg J: **Increased throughput by parallelization of library preparation for massive sequencing.** *PLoS One* 2010, **5**:e10029.

**Citation**

This protocol is derived from the referenced protocol created by Nadin Rohland and David Reich. If you use this protocol, please ensure that you cite their work:

Rohland N, Reich D. **Cost-effective, high-throughput DNA sequencing libraries for multiplexed target capture.** Genome Research 22: 939-946.

**Changes**

- Minor wording changes
- v2.2 – added exact Fisher mix for with-bead library prep
- v2.3 – highlighted ladder issues and added additional warning words for Fisher mix; added additional details about SpeedBeads
- v2.4 – update and cleanup. Add some magnet info.

**Materials**

Here, I list stock solutions that are purchased pre-mixed and sterilized. This is in an attempt to minimize variation to the degree possible. You can certainly prepare your own stock solutions at appropriate pH.

- Sera-mag SpeedBeads (Fisher # 09-981-123; GE # 65152105050250)^[1]^
- PEG-8000 (Amresco 0159)
- 0.5 M EDTA, pH 8.0 (Amresco E177)
- 1.0 M Tris, pH 8.0 (Amresco E199)
- Tween 20 (Amresco 0777)
- 5 M NaCl
- Fermentas ladder(s) (Ultra-low range: Fisher # FERSM1211, 50 bp: FERSM0371)
- Rare-earth magnet stand (Ambion AM10055 or NEB S1506S)

*Optional*

- Agencourt SPRIPlate Super Magnet Plate (Beckman Coulter A32782)
- DynaMag™-96 Side Magnet (ThermoFisher 12331D)

**Steps**

1. In a 50 mL conical using sterile stock solutions, prepare TE (10 mM Tris-HCl, 1 mM EDTA = 500 µL 1 M Tris pH8 + 100 µL 0.5 M EDTA, fill conical to 50 mL mark with dH_2_0).
2. Mix Sera-mag SpeedBeads and transfer 1 mL to a 1.5 mL microtube.
3. Place SpeedBeads on magnet stand until beads are drawn to magnet.
4. Remove supernatant with P200 or P1000 pipetter.
5. Add 1 mL TE to beads, remove from magnet, mix, return to magnet.
6. Remove supernatant with P200 or P1000 pipetter.
7. Add 1 mL TE to beads, remove from magnet, mix, return to magnet.
8. Remove supernatant with P200 or P1000 pipetter.
9. Add 1 mL TE to beads and remove from magnet. Fully resuspend and set microtube in rack (i.e. not on magnet stand).
10. Add 9 g PEG-8000 to a new 50 mL, sterile conical.
11. Add 10 mL 5 M NaCL (or 2.92 g) to conical.
12. Add 500 µL 1 M Tris-HCL to conical.
13. Add 100 µL 0.5 M EDTA to conical.
14. Fill conical to ~ 49 mL using sterile dH_2_0. You can do this by eye, just go slowly.
15. Mix conical for about 3-5 minutes until PEG goes into solution (solution, upon sitting, should be clear).
16. Add 27.5 µL Tween 20 to conical and mix gently.
17. Mix 1 mL SpeedBead + TE solution and transfer to 50 mL conical.
18. Fill conical to 50 mL mark with dH_2_0 (if not already there) and gently mix 50 mL conical until brown.
19. Test against AMPure XP using aliquots of ladder (Fermentas GeneRuler). I recommend the 50 bp ladder in place of the ultra-low range ladder.
20. Wrap in tinfoil (or place in dark container) and store at 4 °C.
21. Test monthly – see Testing, next page.

You may also wish to prep an extra 50 mL of PEG solution that lacks Sera-mag SpeedBeads so that you can use it in a bead-inclusive library preparation protocol. You can either use the recipe from above, but leave out the beads & fill to 50 mL or you can use the recipe below, derived from Fisher (2011):

1. Add 10 g PEG-8000 to a new 50 mL, sterile conical.
2. Add 25 mL 5 M NaCL (or 7.3 g) to conical.
3. Fill conical to ~ 49 mL using sterile dH_2_0. You can do this by eye, just go slowly.
4. Mix conical for about 3-5 minutes until PEG goes into solution (solution, upon sitting, should be clear).
5. Fill conical to 50 mL mark.

**Note:** this has a slightly higher concentration of PEG (20% vs. 18%) & much higher concentration of NaCl (2.5 M vs. 1M), which probably slightly changes the size-range of fragments recovered (though we tend to use them interchangeably).

**Testing**

You should test the SpeedBead mixture to ensure that it is working as expected. You can do this using DNA ladder and we suggest Fermentas GeneRuler – because many other ladders do not work appropriately):

1. Prep fresh aliquots of 70% EtOH.
2. Mix 2 µL GeneRuler with 18 µL dH_2_0.
3. Add 20 µL GeneRuler mixture to a volume of SpeedBead and/or AMPure (the specific volume depends on whether you are trying exclude small fragments or not; see the figure on the next page).
4. Incubate mixture 5 min. at room temperature.
5. Place on magnet stand.
6. Remove supernatant.
7. Add 500 µL 70 % EtOH.
8. Incubate on stand for 1 min.
9. Remove supernatant.
10. Add 500 µL 70% EtOH.
11. Incubate on stand for 1 min.
12. Remove supernatant.
13. Place beads on 37°C heat block for 3-4 min. until dry.
14. Rehydrate with 20 µL dH_2_0.
15. Place on magnet stand.
16. Transfer supernatant to new tube.
17. Mix supernatant with 1 µL loading dye.
18. Electrophorese in 1.5 % agarose for 60 min. at 100 V.

The following image compares the results of “purifying” a mix of 2 µL Fermentas Ultra Low Range Ladder + 18 µL dH_2_0 using several different amounts of AMPure or SpeedBead solution to DNA solution. AMPure is on the left, “SpeedBead” is on the right. After preparing 20 µL of ladder + water mix, we combined that with the volumes of AMPure or SpeedBead listed below and then purified using the standard protocol:

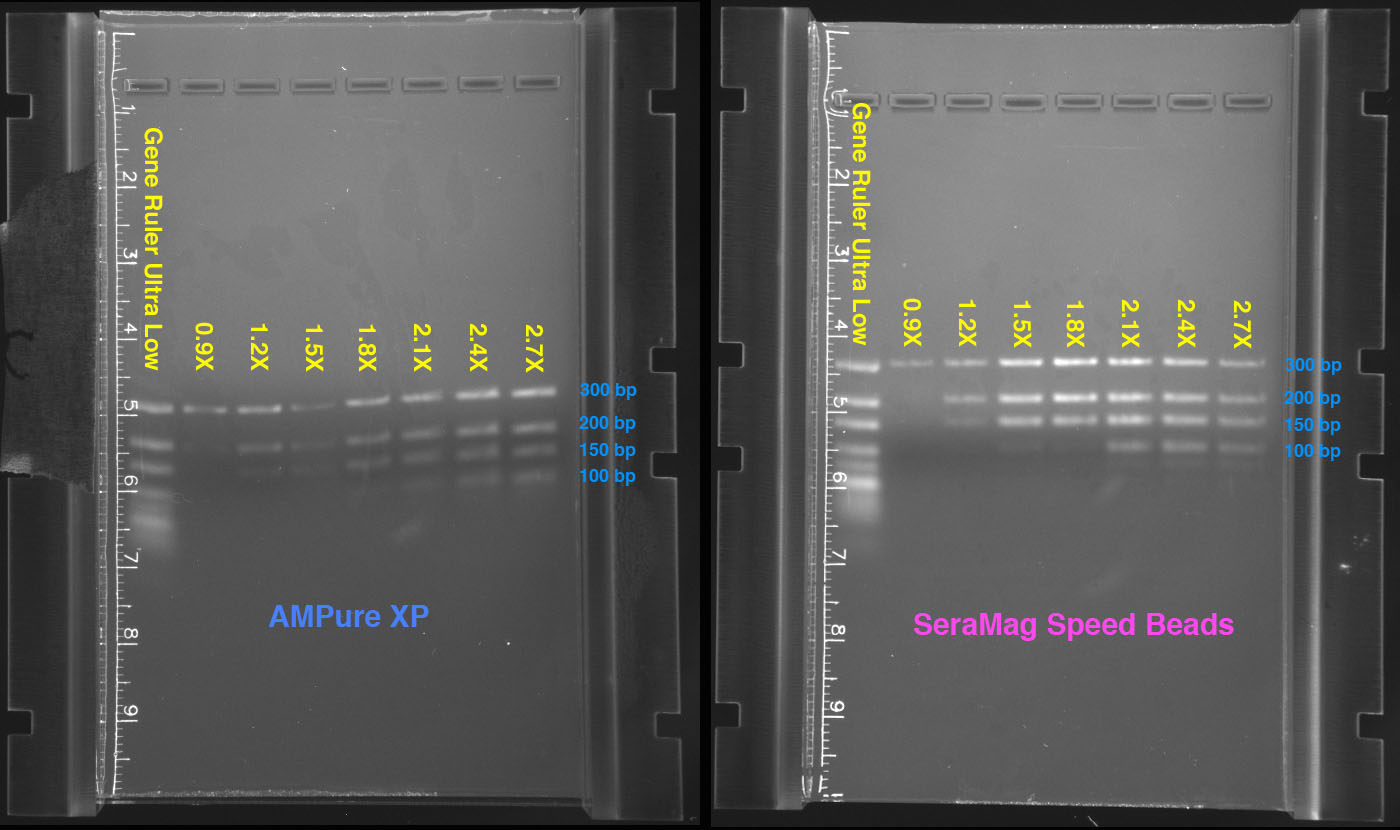

As you can see, the volume of AMPure or SpeedBead controls the size of fragments recovered. More specifically, it is the ratio of PEG solution used to the volume of the DNA in solution which makes the difference, not the count of beads in solution (provided they are above the minimum level). This is what makes it possible to do “double-SPRI” size selection.

1. These are Sera-Mag SpeedBead Carboxylate-Modified Magnetic Particles (Hydrophobic), 15 mL ~$375; the same beads used by Beckman, per Orapure product sheet and <http://bit.ly/vmiDzU>)
