## Supplementary material for "Adapterama I: Universal stubs and primers for 384 unique dual-indexed or 147,456 combinatorially-indexed Illumina libraries (iTru & iNext)": File S9

**Overview:** We are simply following the methods for Kapa Hyper Prep library preparation kits, but using half volumes and Speedbeads.

**Protocol:**

1. Normalize DNA at ~23 ng/µL
2. Bioruptor fragmentation: 3 sets of 5 cycles (15 cycles) on Medium setting.
3. Insert 25 µL of sheared DNA in 10 µL of End Repair Master Mix.
4. Incubate in Thermocycler at 20 **°**C for 30 minutes.
5. Add 100 µL of SeraMag speedbeads to 35 µL volume, mix and let sit for 5-15 minutes.
   - Magnet capture & EtOH wash twice with 80 µL of 80% EtOH.
6. To dry beads, add 25 µL of A-tailing master mix.
7. Incubate in Thermocycler at 30 **°**C for 30 minutes.
8. Add 45 µL of PEG/NaCl to 25 µL volume, mix and let sit for 5-15 minutes.
   - Magnet capture & EtOH wash twice with 80 µL of 80% EtOH.
9. To dry beads, add 22.5 µL of Ligation master mix + 2.5 µL of 5 µM iTru Y-yoke adapters.
10. Incubate in Thermocycler at 20 **°**C for 15 minutes.
11. Add 25 µL of PEG/NaCl to 25 µL volume, mix and let sit for 5-15 minutes.
    - Magnet capture & EtOH wash twice with 80 µL of 80% EtOH.
12. To dry beads add 100 µL of TLE (or IDTE; 10 mM Tris pH8, 0.1 mM EDTA).
13. Add 55 µL of PEG/NaCl to 100 µL volume, mix and let sit for 5-15 minutes.
14. Magnet capture, transfer clear supernatant (~155 µL) to new tube and discard the old speedbeads.
15. Add 25 µL of speedbeads to 155 µL volume of transferred supernatant, mix and let sit for 5-15 minutes.
    - Magnet capture & EtOH wash twice with 80 µL of 80% EtOH.
16. To dry beads add 25 µL of TLE (or IDTE). This is the Pre-PCR Library Prep.
17. PCR: To a new tube add 12.5 µL of KAPA Hotstart Ready Mix, 1.25 µL of i5 primer, 1.25 µL of i7 primer, and 10 µL of the Pre-PCR Library Prep with speedbeads, mix and place in thermocycler for 8-12 cycles.
18. Run 5 µL on a gel.
19. For samples that need it, amplify with more cycles.
20. Cleanup: add 25 µL of speedbeads to 20 µL volume.
21. Quantify on Qubit.
