## Supplementary material for "Adapterama I: Universal stubs and primers for 384 unique dual-indexed or 147,456 combinatorially-indexed Illumina libraries (iTru & iNext)": File S15: countIndexes.py

```
#!/usr/bin/env python
# -*- coding: utf-8 -*-
 
"""
(c) 2014 Brant Faircloth || http://faircloth-lab.org/
All rights reserved.
 
This code is distributed under a 3-clause BSD license. Please see
LICENSE.txt for more information.
 
Created on 14 October 2014 09:35 CDT (-0500)

print commands & additional comments for idiots added by TCG in November 2014.

This program assumes that you have an undetermined.fasta.gz file with dual indexes
You need to put this file wherever your IDLE points (default = documents)

I strongly suggest that you use the UndeterminedTestFile.fasta.gz to ensure all is well 
before searching a big file!

In this program, you will need to change the input file name to whatever is appropriate.  
The input fileName is currently: countMyIndexes.fastq.gz
That inputFileName is on line 43 (IMPORTANT) & then copied to line 38 so you know the filename you used
The print function of line 38 confirms that your script is running (or at least started correctly)

The  UndeterminedTestFile.fasta.gz runs in just a few seconds.  
A real file (3 GB, 30M reads) takes 20 minutes to run (your mileage may vary).

I have this set to print the 50 most common indexes, but you can change that to any number you want
"""
 
import gzip
from collections import Counter
from pprint import pprint

#this is the input filename - COPIED from 7 lines below (in the line that starts: with gzip.open...)
print
print "You are now searching the file countMyIndexes.fastq.gz"
print

#Change the Undetermined fileName 2 lines below this line!!!
barcode_combo_count = Counter()
with gzip.open('countMyIndexes.fastq.gz', 'rb') as infile:
    for line in infile:
        if line.startswith('@'):
            ls =line.strip().split(' ')
            barcodes = ls[-1]
            barcode_combo_count.update([barcodes.split(":")[-1]])
 
 
# get most common 10 [commented out by TCG because we're printing these below]
#barcode_combo_count.most_common(10)

# get total count of all reads [commented out by TCG because we're printing these below]
#sum(barcode_combo_count.values())

# get a subset of reads matching some barcode pattern [commented out by TCG because I'm not using this option]
# assumes dual-indexes
# new_counter = Counter({i:j for i,j in barcode_combo_count.iteritems() if "+AGATCTCG" in i})

#More TCG comments and output for idiots --

#print the total count of all reads - comment the next line if you don't want to see it

print "Here is the sum of all sequences in the file with indexes:"
print sum(barcode_combo_count.values())
print

#print the most common 50 barcodes - comment the next line if you don't want to see it
print "Here are the 50 most common indexes in the file:"
print
pprint (barcode_combo_count.most_common(50))
print
```
